# Active bulging promotes biofilm formation in a bacterial swarm

**DOI:** 10.1101/2022.08.25.500950

**Authors:** Siyu Liu, Ye Li, Haoran Xu, Daniel B. Kearns, Yilin Wu

**Affiliations:** Department of Physics and Shenzhen Research Institute, The Chinese University of Hong Kong, Shatin, NT, Hong Kong, P.R. China; Department of Biology, Indiana University, Bloomington IN, USA

**Author notes:** Correspondence: Yilin Wu. These authors contributed equally (Siyu Liu, Ye Li).

## Abstract

Microbial communities such as biofilms are commonly found at interfaces. However, it is unclear how the physical environment of interfaces may contribute to the development and behavior of surface-associated microbial communities. Combining multi-mode imaging, single-cell tracking and numerical simulations, here we discovered that an interfacial process denoted as “active bulging” promotes biofilm formation. During this process, an initially two-dimensional layer of swarming bacteria spontaneously develops scattered liquid bulges; the bulges have a higher propensity to transit from motile to sessile biofilm state, presumably due to the enrichment of pre-existing immotile cells in the colony. We further demonstrate that the formation of liquid bulges can be controlled reversibly by manipulating the speed and local density of cells with light. Our findings reveal a unique physical mechanism of biofilm formation and provide a new strategy for biofilm patterning in engineered living materials as well as for directed self-assembly in active fluids.

## Introduction

Microbial communities of ecological and clinical importance are commonly found in interface-associated environments, such as biofilms (1, 2) and bacterial swarms (3, 4) developed on solid substrates, pellicles formed at liquid-air interface (5), and microbiome thriving in the gastrological tract of animals (6). In the past two decades, much has been learned on the development and behavior of surface-associated microbial communities in terms of their genetic programs (7-10) and chemical communications (such as quorum sensing (11) and vesicles delivery (12, 13)). More recently, mechanical forces at the interfaces are expected to play crucial roles in the surface-associated life of microbial communities (14-17). For instance, cell-substrate mechanical interaction drives biofilm morphogenesis and orientational ordering of cells (18, 19); while surface flows have been shown to promote the expansion (20-25) and material transport (26) within bacterial colonies. However, it remains largely unexplored how the physical environment of interfaces may contribute to more diverse processes involved in the development and behavior of surface-associated microbial communities.

Here we focus on the transition between the motile and sessile state of bacterial communities grown on solid substrates, a key step in the adaptation to environmental fluctuations of bacterial communities (16, 27, 28). During the transition, cells switch their gene regulation program from expressing flagellar genes to one that expresses biofilm genes and deactivates flagellar motility (29). The transition generally requires surface-sensing mediated by cell surface appendages (30-35) or envelope stress (36, 37), but can also be facilitated by biophysical mechanisms such as motility-induced phase separation and motility segregation (38, 39). In this study we discover that a novel interfacial process denoted as “active bulging” promotes the transition from motile to sessile biofilm state of bacterial communities. During the process, an initially two-dimensional layer of swarming motile cells spontaneously develops scattered liquid bulges manifesting as regions where cells are highly motile and can pile on top of one another. The liquid bulges maintain a constant difference (∼20% higher) in surface-packing cell density from elsewhere; as time evolves, the liquid bulges have a higher propensity to transit from motile to sessile or biofilm state. Using a multi-mode imaging system, we show that the formation of liquid bulges can be controlled reversibly by manipulating the speed and local density of cells, which is explained by an active Brownian dynamics model. Our experiments and simulations further suggest that immotile cells (i.e., cells with deactivated flagellar motility) pre-existing in the colony continuously get enriched in the liquid bulges due to differential diffusivity inside and outside the bulges, thus increasing the propensity of biofilm formation in the liquid bulges. The findings illustrate the importance of interfacial mechanics to a comprehensive understanding of bacterial behavior and dynamics. Our results also provide a new strategy to control biofilm patterns in engineered living materials (40-47) and for directed self-assembly in active matter fluids (48-51).

## Results

### Liquid bulges spontaneously emerge in homogeneous *B. subtilis* swarms

To study motile-to-sessile state transition in surface-associated bacterial communities, we chose to work with bacterial swarms (3, 4). Bacterial swarming resembles the early stage of biofilm formation on solid substrates (27), during which bacterial colonies extract liquid from the surroundings to form a free-surface liquid film that covers the entire colony and supports flagellar motility (52, 53). *Bacillus subtilis* is a model microorganism that displays robust swarming behavior (4). Here, we found that after extended growth, an initially homogeneous two-dimensional monolayer of motile cells in *B. subtilis* (wild type NCIB3610) swarms spontaneously develops a pattern of scattered domains (Fig. 1A; Movie S1; Methods) with higher surface-packing density of cells (Fig. 1B; Methods). The boundaries of such domains are dynamic but well-defined in low magnification phase-contrast images (Fig. 1A) or in darkfield microscopy images (Fig. 1B). Cells within the high-surface-density domains move freely with the mean speed and persistence time only slightly lower (∼7% for mean speed and ∼9% for persistence time) than elsewhere (Fig. 1C,D; Methods), which is different from the clustering domains formed in *B. subtilis* swarms via motility-induced phase separation (38, 54) that largely consist of jammed cells., In contrast to cells moving in a strictly two-dimensional (2D) manner (i.e. as a monolayer) outside these domains, cells inside the domains often overlap with each other (Movie S2), suggesting that the height of swarm fluid film in these domains is higher than in the monolayered regions. Therefore these domains with higher surface-packing density are in fact bulges of the swarm fluid film at its free surface where more motile cells can be accommodated per unit surface area.

**Fig. 1.**
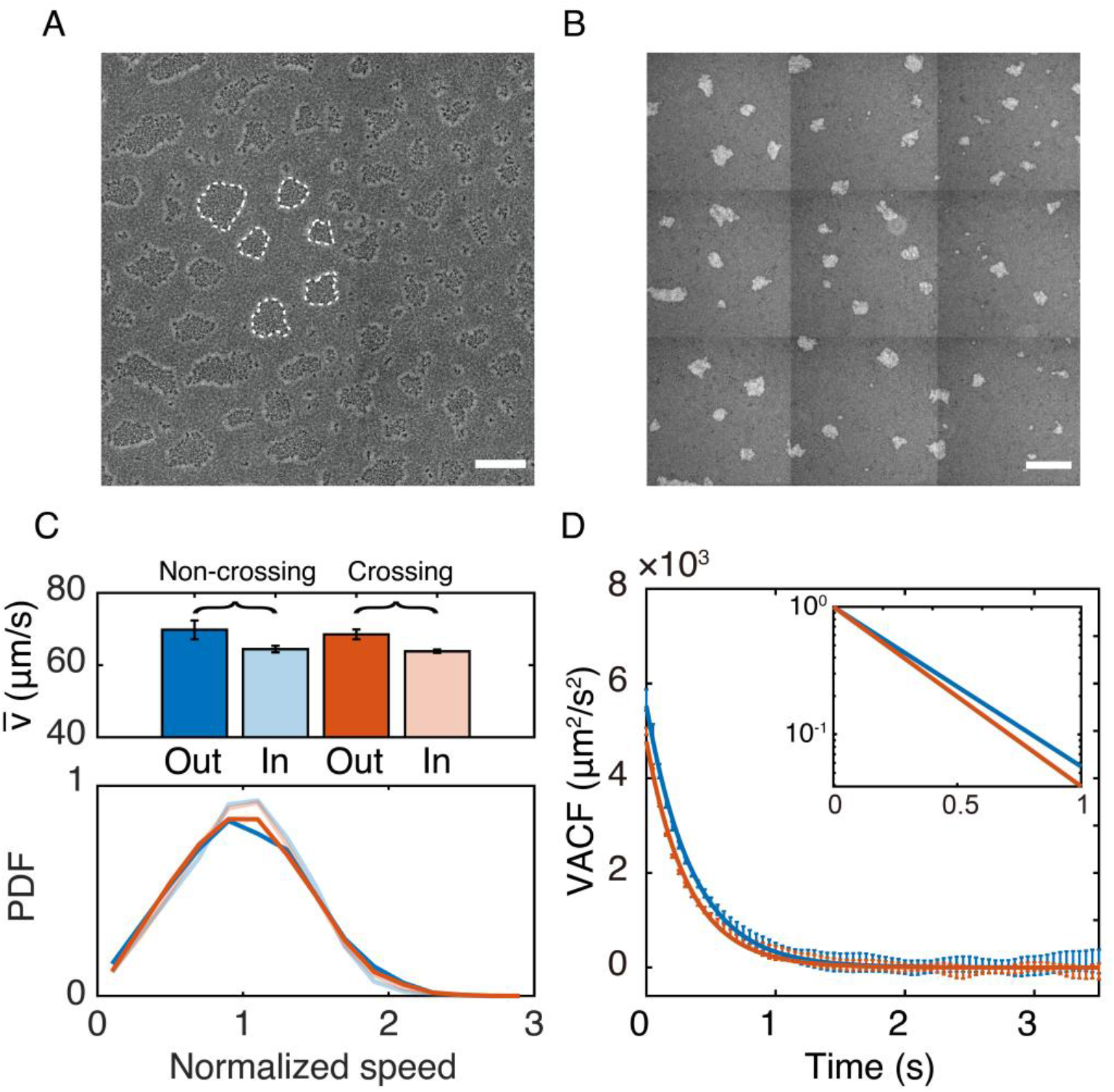
Liquid bulges with higher surface-packing cell density emerge in *B. subtilis* swarms. (A) A representative large-field phase-contrast microscopy image (Methods) showing a pattern of scattered domains in the swarming colonies of *B*.*subtilis*. Dash lines indicate the boundaries of such domains. (B) A representative large-field darkfield microscopy image of the scattered-domain pattern similar to that shown in panel A (Methods). In dark field microscopy, the brightness is positively correlated with surface-packing density of cells, thus brighter domains in the image have higher surface-packing cell density. Panels A,B were obtained by stitching overlapping images taken in a smaller field of view. Scale bars in panels A,B, 500 μm. (C) Motion pattern of cells inside and outside the high-surface-density domains or liquid bulges. Upper panel: Average speed 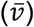 of cells that remained outside and inside the high-surface-density domains during the tracking period are shown as dark blue (69.8 ± 2.6 μm/s; mean ± SE, N = 5) and light blue (64.5 ± 0.9 μm/s; mean ± SE, N = 5) columns, respectively; the average speed of cells that crossed the boundaries of the domains during the tacking period are shown as dark red (while moving outside the domains; 68.5 ± 1.3 μm/s, mean ± SE, N = 4) and light red (while moving outside the domains; 63.8 ± 0.5 μm/s, mean ± SE, N = 4) columns, respectively. Lower panel: Average probability density functions (PDFs) of the normalized speed of cells corresponding to the 4 scenarios in the upper panel, following the same color coding as that of the columns in upper panel. The speed was normalized by the mean speed of all cells in the corresponding scenario. (D) Velocity autocorrelation functions (VACFs) of cells that remained inside and outside the liquid bulges computed based on single cell trajectories. Solid lines are exponential fits to the VACFs (blue: outside liquid bulges, VACF_out_ ∼ exp(-t/0.35), persistence time *τ*_out_ = 0.35 ± 0.01 s; red: inside liquid bulges, VACF_in_ ∼ exp(-t/0.32), persistence time *τ*_in_ = 0.32 ± 0.01 s). The diffusivity computed from VACF is ∼20% lower inside bulges than outside (∼980 μm^2^/s vs. ∼770 μm^2^/s) (Methods). Plotted in the inset are normalized fitted VACFs outside (blue) and inside (red) liquid bulges in semi-logarithmic scale. Error bars represent the standard deviation of N=5 biological replicates.

To examine whether cells inside and outside the liquid bulges represent different sub-populations with intrinsic motility difference, we tracked the motion of cells near the bulge boundaries. We found that cells can freely cross the boundary (Movie S2); the same cell had a lower mean speed inside bulges than outside the bulges (Fig. 1C, left), with magnitudes similar to those that had not crossed the boundaries during the period of tracking. In other words, when cells entered the liquid bulges, their speed decreased slightly and vice versa. These results suggest that the slight change of motion pattern (i.e., differences in speed and persistence time) of cells with identical intrinsic motility is due to the difference in the local physical environment inside liquid bulges. Moreover, the probability density functions (PDFs) of normalized bacterial speed inside and outside liquid bulges are similar (Fig. 1C, right). These results together show that the speed difference inside and outside liquid bulges is due to the difference in local physical environment but not due to the presence of sub-populations with intrinsic motility difference. Nonetheless, immotile cells with deactivated flagellar motility can naturally arise in a swarm due to cell-cell variability of motility (55, 56) or due to the genetic program of motile-to-sessile transition (29, 57-59), and a recent study suggests that immotile subpopulation nucleates stationary clusters in *B. subtilis* swarms (39).

To examine whether immotile cells in the swarm contribute to liquid bulge formation, we chose a Δ*sinI* mutant of *B. subtilis*, which is incapable of differentiation into sessile cells and thus retains active flagellar motility (60, 61). We found a similar phenomenon of liquid bulge formation in swarms of *B. subtilis* Δ*sinI* mutant (Movie S3), and cells inside and outside the liquid bulges displayed similar motion patterns compared to their counterparts in wildtype *B. subtilis* swarms (Fig. S1). These results show that liquid bulge formation does not require an immotile sub-population of cells in the swarm. Taken together, the liquid bulges we report here are different from those patterns of scattered domains found in bacterial colonies whose formation is driven by motility-induced phase separation (38, 54) or by nucleation of immotile cells (39).

### Liquid bulges have a higher propensity to transit into biofilm state

Over time, the liquid bulges grew in size and passively slid along the colony expansion direction; meanwhile, nearby bulges along the colony expansion direction coalesced and fused into stripes (Fig. 2A; Movie S1). For wildtype swarms during this process, cells outside the bulges remained highly motile, but the motility of cells inside the bulges appeared to decrease (Fig. 2B). After ∼2-3 hr from the emergence of liquid bulges, most cells in the bulges became immotile and cells appeared to have a chain-like morphology characteristic of cells in *B. subtilis* biofilms (5) (Fig. 2A, right; Movie S4). By contrast, Δ*sinI*, did not chain and cells remained highly motile everywhere as liquid bulges continued to merge with each other until covering the entire field of view (Movie S3), at which time the swarm became homogeneous again and the height of the swarm fluid film must be uniform. These observations suggest that liquid bulges of wildtype swarms have a higher propensity than elsewhere to transit from motile to sessile or biofilm state. To examine this idea, we monitored the physiological state of wildtype cells inside and outside liquid bulges with three fluorescence reporters starting from the emergence of stable liquid bulges (Methods): P_*tapA*_-*gfp*, a genetically encoded fluorescence reporter for the expression level of *tapA* gene (encoding an anchor protein TapA for biofilm matrix component TasA) indicating the extent of biofilm maturation (62-64); P_*hag*_-*gfp*, a genetically encoded fluorescence reporter for the expression level of *hag* gene (encoding flagellin protein) serving as an indicator of cell motility (55, 62); and the lipophilic dye FM4-64 that stains the cell membrane and indicates cell density (65).

**Fig. 2.**
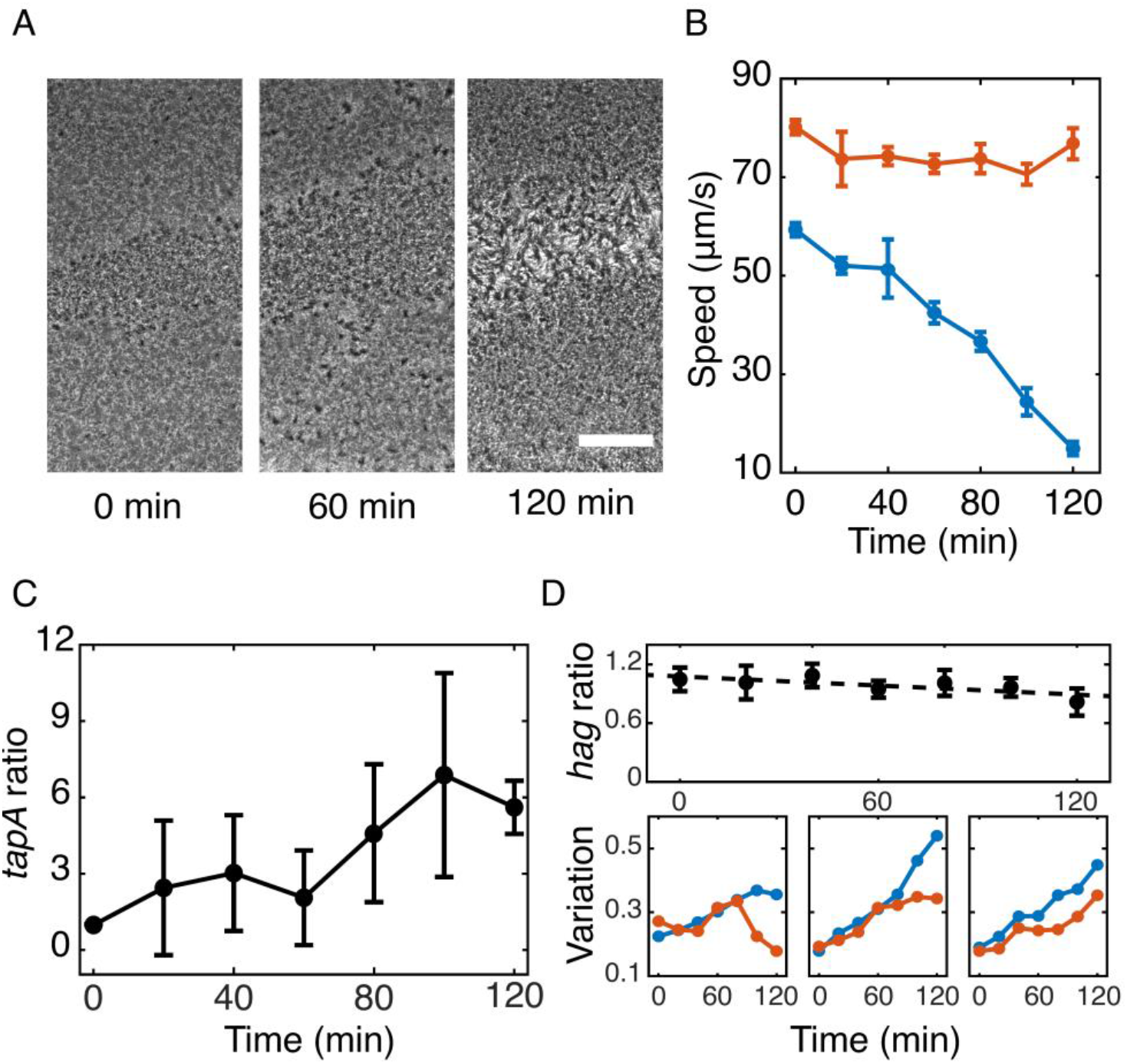
Transition of cells in bulges into biofilm state. (A) Snapshots of phase-contrast microscopy image sequences showing the development of liquid bulges in a wildtype swarm. Striped liquid bulges are shown as patterns at the center of images along the horizontal direction with different contrast. Scale bar, 300 μm. (B) Temporal dynamics of average collective speed of cells inside (blue) and outside (red) liquid bulges in a wildtype swarm computed by optical flow analysis (Methods). Error bars represent standard deviation of collective speed distributions in the two regions. (C) Temporal dynamics of the ratio between average single-cell P_*tapA*_-*gfp* reporter activity inside and outside liquid bulges. Error bars represent standard deviation (N = 3). (D) Temporal dynamics of single-cell P_*hag*_-*gfp* reporter activity. Upper: The ratio between average single-cell P_*hag*_-*gfp* reporter activity inside and outside liquid bulges plotted against time (indicated in the abscissa). Dash line is a least-squares linear fit with slope ∼ -0.0015 (*R*^*2*^ = 0.58). Error bars represent standard deviation (N = 4). Lower: Normalized spatial variation of the P_*hag*_-*gfp* reporter fluorescence inside (blue) and outside (red) bulges (Methods); data in each panel is from one representative experiment and the dynamics show variability across experiments. Solid lines in panels B-D serve as guides to the eye.

We found that the ratio between single-cell P_*tapA*_-*gfp* reporter activities inside and outside liquid bulges gradually increased throughout the developmental process (Fig. 2C). On the other hand, the ratio between single-cell P_*hag*_-*gfp* reporter activities inside and outside liquid bulges only decreased slightly throughout the developmental process (Fig. 2D, upper). This result is unexpected, because the expression levels of *tapA* and *hag* are supposed to be inversely correlated (62) and thus one would expect a substantial decrease in the ratio of P_*hag*_-*gfp* reporter activities. However, close inspection of the fluorescence images revealed that liquid bulges displayed larger spatial variation of the P_*hag*_-*gfp* reporter fluorescence after ∼80 min from their emergence (Fig. 2D, lower), with randomly spaced cellular clusters of much higher P_*hag*_-*gfp* reporter fluorescence than elsewhere (Fig. S2); the brighter fluorescence from these clusters masked the reduced P_*hag*_-*gfp* reporter activity of chained cells inside liquid bulges. Cellular clusters with high P_*hag*_-*gfp* reporter fluorescence likely arose from stochastic switching from chained to motile state (58, 59), and they appeared immobile because of jamming or entrapment in between the chained cells. Overall, the reporter results show that liquid bulges of wildtype swarms indeed are more prone to transiting into the biofilm state. Thus the formation of liquid bulges represents an unrecognized biophysical mechanism that promotes multilayer development preceding biofilm formation and is distinct from mechanisms driven by motility-induced phase separation (38) or by nucleation of immotile cells (39).

### Liquid bulge formation depends on cell motility and density

Next, we sought to understand how the liquid bulges formed and followed the developmental process of the liquid bulges in *B. subtilis* swarms. We found that the formation of liquid bulges was preceded by the spontaneous nucleation of transient domains with high surface-packing density of cells with various size of ∼10-60 μm and ∼1-2 s lifetime (Movie S5). Within several minutes, the transient domains grew in size and gradually stabilized into liquid bulges with well-defined boundaries. The results indicate that the formation of liquid bulges is associated with transient increase in surface-packing density of cells due to density fluctuations in the swarm often seen in self-propelled particle systems driven by local free energy input or particle motility (66, 67).

To understand how cell motility and local cell density variation relate to the formation of liquid bulges, we manipulated cell motility *in situ* in the swarm; here the non-chaining *sinI* mutant was used in order to exclude the artifact in speed measurement brought by any pre-existing immotile cells. Violet light illumination (∼406 nm) was used to reversibly control cell speed because violet light generates reactive oxygen species that suppress flagellar motility (68) (Fig. S3; Methods). When we illuminated a region with pre-existing liquid bulges (Fig. 3A and Movie S6), cell speed decreased from ∼65 μm/s to ∼22 μm/s under violet light illumination, and the local cell number density fluctuation Δ*N*/*N* (Methods) in the illuminated monolayered regions was decreased; meanwhile, the existing liquid bulge gradually dispersed. Withdrawal of violet light illumination allowed cell speed recovery to ∼45 μm/s and new liquid bulges emerged. The result shown in Fig. 3A demonstrated that the degree of local cell density fluctuation driven by cell motility is strongly correlated with bulge formation.

**Fig. 3.**
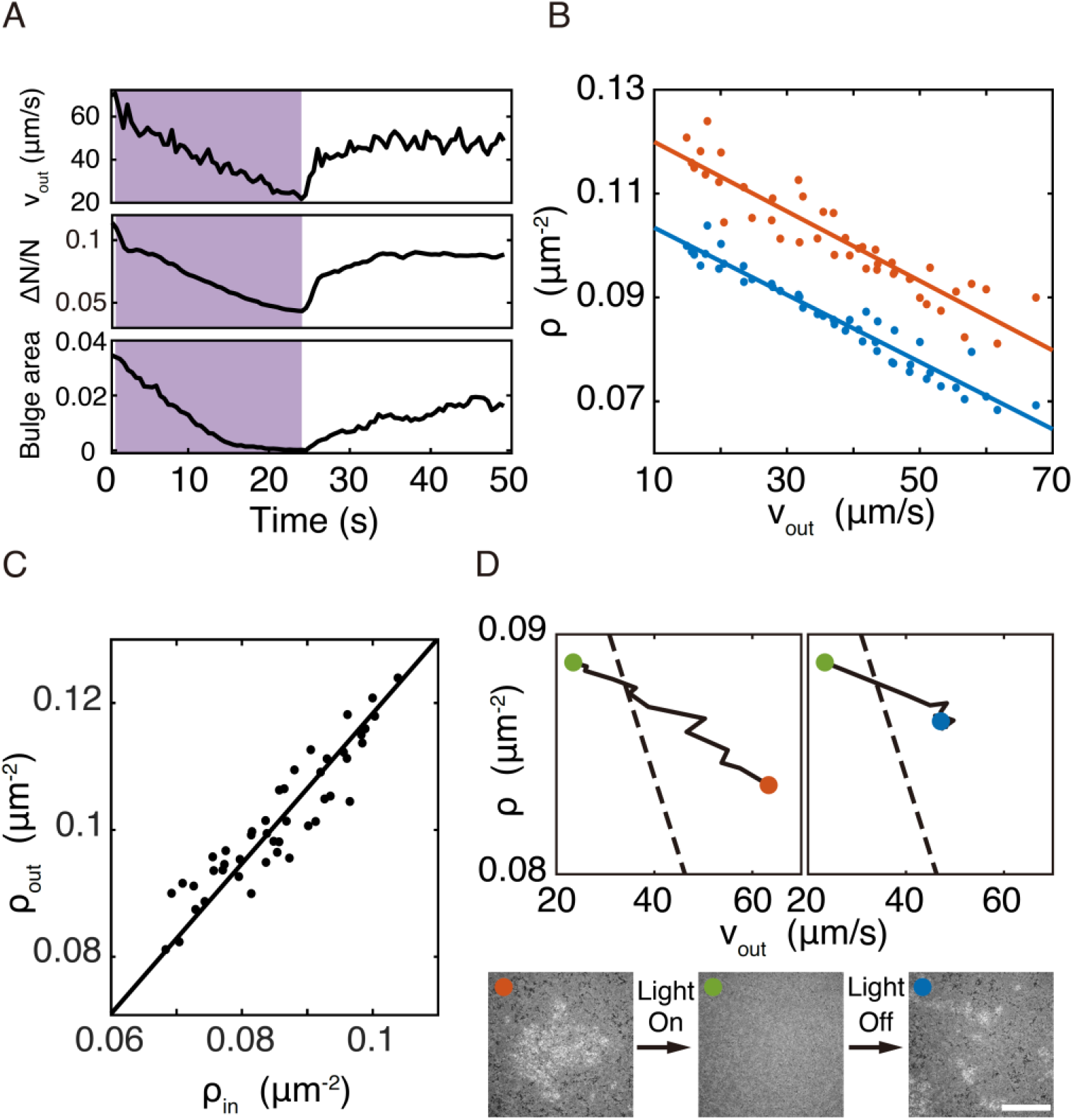
Motility and density dependence of bulge formation in B. subtilis swarms. (A) Local cell density fluctuation is correlated with liquid bulge formation. Plotted here are temporal dynamics of collective cell speed outside bulges obtained by optical flow analysis on dark field microscopy images (*ν*_*out*_; upper), cell density fluctuation (*ΔN*/*N*, where *ΔN* and *N* are the standard deviation and average of cell number, respectively; middle), and area fraction of bulges (lower); see Methods. Violet light was illuminated during the period shaded in purple color. (B) Relation between cell speed and surface-packing cell number density for monolayered regions (blue) and for the coexisting liquid bulges (red) (Methods). Collective cell speed measured in the monolayered regions (*ν*_*out*_; Methods) was used in the abscissa. Lines are least-squares linear fits to the data (*R*^*2*^ = 0.94 for blue line and 0.85 for red line). Data presented here was from 3 biological replicates. (C) Relation between surface-packing cell density inside (*ρ*_*in*_) and outside (*ρ*_*out*_) bulges. Solid line is least-squares linear fit with *ρ*_*in*_ = 1.18*ρ*_*out*_ (*R*^*2*^ = 0.83). Data presented here was from 3 biological replicates. (D) Phase trajectories in the plane of cell speed and density during reversible manipulation of the liquid bulge formation in panel A (also see Movie S6). In the phase trajectories, red dot represents the initial state at the beginning of violet light illumination when liquid bulges were present (Time = 0.3 s); green dot represents the state at the end of violet light illumination, at which time liquid bulges had completely dispersed (Time = 24.3 s); blue dot represents the state when bulges re-formed due to motility recovery (Time = 49.8 s). Darkfield microscopy images corresponding to the three time points are shown below the phase trajectory plots labeled in the same marker (scale bar, 300 μm). Dash lines are a portion of the fitted speed-density relation for monolayered regions that coexist with liquid bulges as obtained in panel B.

Nonetheless, we noted that the average surface-packing density of cells in a monolayered area illuminated by violet light would become higher than that outside the illuminated area (Fig. S4), presumably due to reduced cellular flux out of the illuminated area during speed decrease, a phenomenon predicted in self-propelled particles (69) and observed in active bacterial suspensions (38, 50, 51, 70). For instance, in the experiment shown in Fig. 3A, the average surface-packing density of cells in the illuminated monolayered regions was increased by ∼10% (Fig. S4), close to the level of cell density fluctuations at normal speed (∼10%) and a significant magnitude compared to the maximal transient increase of local cell density due to the fluctuations (∼20%). Why wouldn’t the violet light-induced increase in cell density compensate for the decreased level of local cell density fluctuation and thus prevent the dispersal of liquid bulges during cell speed reduction? This question prompted us to investigate the detailed relation between cell motility and surface-packing density of cells inside and outside liquid bulges. For this purpose, we designed a multi-mode imaging system that integrates large-scale (∼3 cm^2^) blue/violet light illumination (for speed reduction in the swarm), darkfield microscopy (for speed measurement by optical flow analysis), and wide-field fluorescence microscopy (for surface-packing density measurement by counting the number of fluorescently labeled non-chaining cells) (Fig. S5; Methods).

Using the multi-mode imaging system, we found that the surface-packing density of cells showed a negative quasi-linear relation with cell speed during bulge formation (Fig. 3B). Interestingly, the ratio between surface-packing densities (i.e., cell number per unit surface area but not per fluid volume) inside and outside liquid bulges remained nearly constant at ∼1.2 regardless of the cell speed (Fig. 3B,C). The similar persistence time of cells inside and outside bulges (as revealed by the velocity autocorrelation shown in Fig. 1C,D) suggests that the volume density of cells is nearly homogeneous across the swarm fluid film; thus the constant ratio of surface-packing densities of cells measured here suggest that the height difference of liquid bulges compared to the monolayered regions remained constant, as long as cell motility is sufficiently high to support bulge formation.

We hypothesized that the quasi-linear speed-density relation measured in monolayered regions that coexist with liquid bulges (blue line in Fig. 3B) represents the motility (or density) threshold at a specific cell density (or cell motility) that would give rise to liquid bulges. Indeed, the experiment in Fig. 3A corresponds to a case that the cell speed crossing this threshold to a lower level that does not support bulge formation, followed by speed increase again crossing the threshold that supports bulge formation again (Fig. 3D). Furthermore, by increasing surface-packing density of cells from ∼0.07μm^-2^ to ∼0.09 μm^-2^ in a monolayered region without bulges via violet light illumination, we found that a bulge emerged after withdrawal of violet light (Movie S7) and it survived for >5 minutes. The motion pattern of cells inside and outside bulges formed this way was similar to that associated with naturally developed liquid bulges (Fig. S6). These results demonstrate the feasibility of reversible manipulation of self-assembled liquid bulges in bacterial swarms guided by the quantitative insight from the speed-density relation.

### Active Brownian dynamics simulation explains the motility and density dependence of bulge formation in terms of activity-induced pressure

The experiments above suggest a biophysical origin of the formation of liquid bulges. Swarm cells entrain a certain amount of liquid in the boundary layer associated with their surfaces; when local cell density is increased, their entrained water will be brought together. An increase in water volume would cause local height increase of the swarm fluid film, thus increasing the interfacial area as well as the interfacial free energy (defined as surface tension multiplied by the interfacial area (71)). If cells are immotile and brought closer due to thermal fluctuations, the local height of liquid film would return to normal in order to minimize interfacial free energy; but for actively moving cells that exert active stresses (72) to the fluid film, the activity-induced pressure of cells (i.e., the force exerted on the fluid interface per unit area) would be able to sustain the interfacial area increase, thus supporting liquid bulge formation.

To further understand the relationship between cell motility and surface-packing density of cells, we computationally modeled bacterial swarming dynamics in the framework of active Brownian particle models (73). Briefly, cells in the model are represented as self-propelled hard rods moving in a quasi-two-dimensional space, whose center positions and orientations follow overdamped Langevin dynamics. The motion of rod-like particles or cells is locally coupled due to intercellular steric interactions. Using this model, we first sought to find out how the activity-induced pressure depends on cells’ active speed (denoted as *ν*_0_) and the surface-packing density of cells (denoted as *ρ*_0_) in the simulation of cell monolayers (Fig. 4A; Methods). Starting from a steady state, the simulations revealed that either increasing the active speed or increasing the cell number density would result in an increase of the activity-induced pressure in a monotonic manner. On the *ν*_0_ *−ρ*_0_ plane we can identify the points or sets of (*ν*_0_, *ρ*_0_) that produce the same activity-induced pressure, and the locus of these points form a contour plot of equal pressure; the contour plots for different activity-induced pressures are shown in Fig. 4A.

**Fig. 4.**
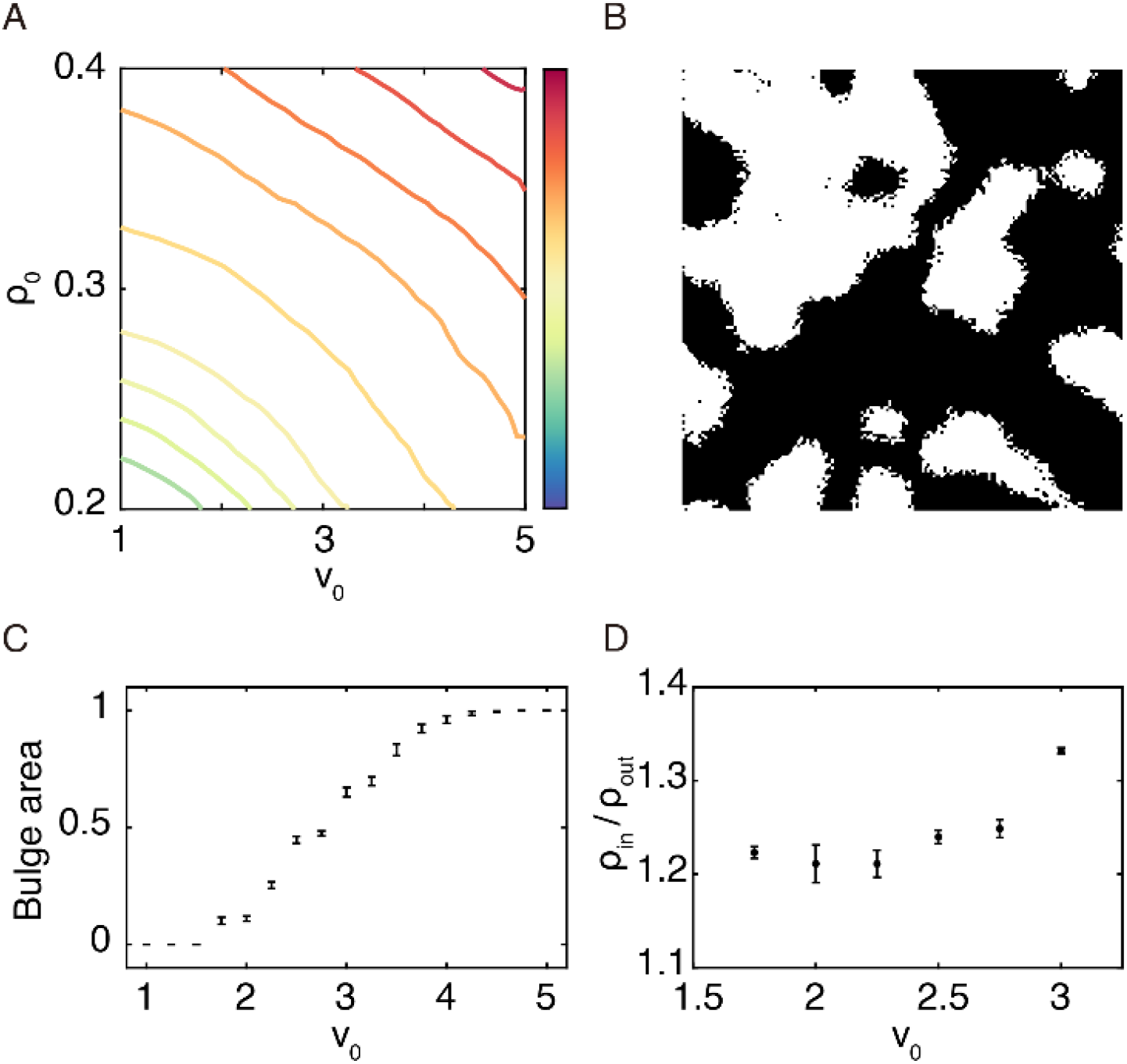
Active Brownian dynamics simulation of bulge formation. (A) Contour lines of activity-induced pressure on the plane of particle active speed (*ν*_0_) and particle number density (*ρ*_0_; unit: particle number per unit area). Colormap to the right indicates the normalized magnitude of activity-induced pressure. (B) Snapshot of bulge formation in a simulation at intermediate active speed *ν*_0_ =2.5. The bright regions denote the bulge area (with lower steric interaction strength; Methods) that persistently coexist with monolayered areas (dark regions). Also see Movie S8. (C) Area fraction of bulges as a function of active speed *ν*_0_ ranging from 1 to 5. Error bars represent standard deviation (N=10 simulation runs). (D) Ratio between the surface-packing cell densities inside (*ρ*_*in*_) and outside (*ρ*_*out*_) bulge area at an intermediate range of *ν*_0_. Error bars represent standard deviation (N=10 simulation runs).

Next, we assume that when local activity-induced pressure is beyond a certain threshold that depends solely on the interfacial properties, the local fluid height will increase (i.e., forming liquid bulge) and thus cells will be able to overlap with each other. This assumption is translated to a lower steric interaction strength in the bulge area in the model (Methods); such a phenomenological treatment avoids the complication of modeling the interaction between cells and the deformable liquid-air interface (74-76), since the height difference of liquid bulges compared to the monolayered regions is expected to remain constant. For a chosen bulge-forming threshold of activity-induced pressure *π*_0_, its contour plot in Fig. 4A allows us to identify the threshold of the local surface-packing density of cells *ρ*_*c*_ for liquid bulge formation at a specific *ν*_0_. By varying *ν*_0_, we found that an intermediate range of *ν*_0_ supports the formation of bulge area (i.e. the region with lower steric interaction strength in the model) that persistently coexist with monolayered regions (Fig. 4B; Movie S8). With the total cell number held constant, we found that the fraction of bulge area increased with cell motility (Fig. 4C; Movie S8), similar to the experimental phenomenon observed during cell speed increase after withdrawal of violet light (Fig. 3A). Moreover, the local surface-packing density of cells is higher inside the bulge area than outside, with the ratio between the surface-packing densities inside and outside the bulge area remaining nearly constant regardless of v_0_ (Fig. 4D), again in agreement with experimental results (Fig. 1C; Fig. 3C).

The simulation results presented above provide a mechanistic explanation for the experiments described above on the motility and density control of bulge formation: Noting that the activity-induced pressure contour plots have a similar shape to the speed-density relation obtained experimentally in monolayered regions that coexist with liquid bulges (see Fig. 3B), the measured speed-density relation resembles the contour line of activity-induced pressure in the experimental system; any deviation to the right from this line (i.e., increasing motility or increasing cell density) would increase the pressure and based on our assumption that liquid bulge formation initiates when local activity-induced pressure is beyond a certain threshold, the local fluid height will increase and lead to liquid bulge formation.

### Expedited biofilm formation in bulges is a result of the enrichment of immotile cells

Finally, we sought to understand why liquid bulges are more prone to transiting into a biofilm state. Immotile cells with deactivated flagellar motility naturally arise in a swarm due to cell-cell variability of motility (55, 56) or due to the genetic program of motile-to-sessile transition (29, 58, 59). With ∼20% higher surface-packing cell density, liquid bulges are expected to contain higher density of pre-existing immotile cells that have naturally arisen in a swarm, and thus liquid bulges would have a higher probability of biofilm transition per unit area. To examine this idea, we would need to track the population distribution of immotile cells inside and out of liquid bulges. However, we lack appropriate reporters of immotile cells that are sufficiently bright for single-cell tracking or population density quantification in swarms (note that the biofilm state reporter P_*tapA*_-*gfp* we used above is very dim and can only detect chained cells that are already in a biofilm state). Instead, we use passive microspheres of a size comparable to cells (diameter ∼1μm) to imitate immotile cells in the swarm and track the motion and spatial distribution of these microspheres (Methods).

We deposited passive microspheres in a lawn of the swarm prior to liquid bulge formation and then followed the position of these microspheres when liquid bulges emerged. Unexpectedly, we found that the density of passive microspheres inside liquid bulges kept increasing (Fig. 5A, black line; Movie S9); meanwhile, the density outside liquid bulges was slowly decreasing (Fig. 5A, red line). This result shows that the ratio between microsphere densities inside and outside bulges is not constant but keeps increasing, which is in stark contrast to the constant ratio of overall surface-packing densities of cells. To explain the enrichment of passive microspheres inside the bulges, we measured the mean speed of microspheres and found that the speed inside bulges is ∼20% lower than outside (∼37 μm/s vs. ∼46 μm/s) (Fig. 5B); meanwhile, the persistence time of microspheres calculated from velocity autocorrelation functions is ∼33% lower (∼0.08 s vs. ∼0.12 s) (Fig. 5C). As a result, the diffusivity of microspheres is ∼53% lower inside bulges than outside (∼84 μm^2^/s vs. ∼178 μm^2^/s) (Methods). Using the computational model described above, we found a similar difference in surface-packing density (Fig. 5D) and a ∼20% difference in diffusivity between immotile cells inside and outside the bulge area. These results can be understood as follows: The diffusivity difference results in an unbalanced particle flux into and out of liquid bulges, which in turn would lead to a persistent increase in the surface-packing density of passive particles in liquid bulges until the particle fluxes are balanced. Taken together, our results based on the motion pattern analysis of passive particles in both experiments and simulations suggest that pre-existing immotile cells tend to get enriched in liquid bulges, which increases the propensity of biofilm development in liquid bulges. Moreover, as the progenies of immotile cells would be less diffusive in liquid bulges than elsewhere, the enrichment of immotile cells may also trigger a positive feedback that expedites biofilm development.

**Fig. 5.**
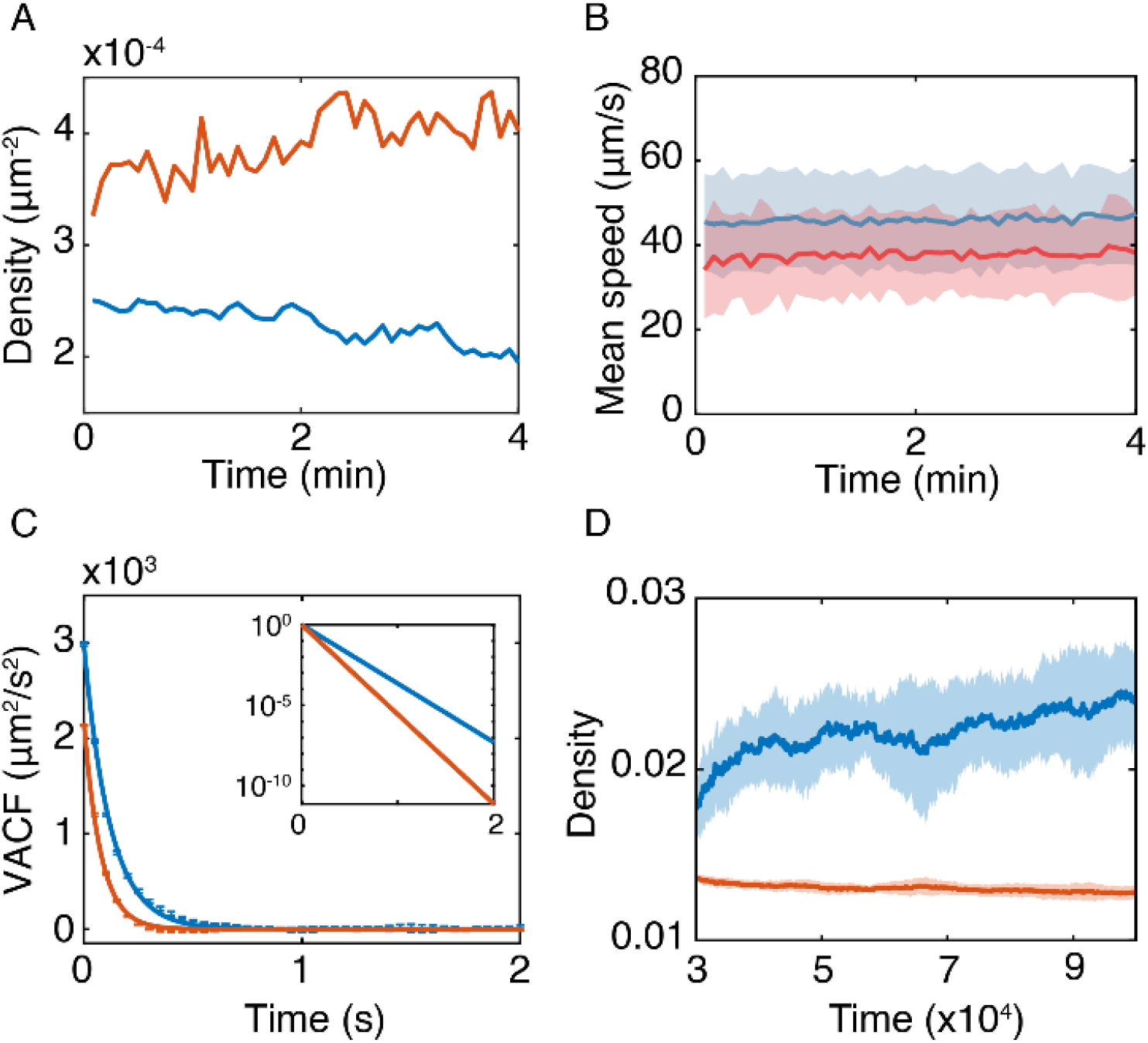
Continuous enrichment of passive particles inside liquid bulges. (A) Surface-packing density of microspheres in bulge area (red) and monolayered regions (blue) plotted against time in a *B. subtilis* swarm. (B) Mean speed of microspheres inside (red line) and outside (blue line) liquid bulges plotted against time. The shaded area around each line represents the standard deviation of microsphere speed inside (red) and outside (blue) liquid bulges. Data in panels A,B was from a representative experiment (3 biological replicates). (C) Velocity autocorrelation functions (VACFs) of microsphere that remained inside and outside the liquid bulges computed based on single-particle trajectories. Solid lines are exponential fits to the VACFs [blue: outside liquid bulges, VACF_out_ ∼ exp(-t/0.12), persistence time *τ*_out_ = 0.12 ± 0.01 s; red: inside liquid bulges, VACF_in_ ∼ exp(-t/0.08), persistence time *τ*_in_ = 0.08 ± 0.01 s). Plotted in the inset are normalized fitted VACFs outside (blue) and inside (red) liquid bulges in a semi-logarithmic scale. Error bars represent the standard deviation of N=3 biological replicates. (D) Surface-packing density of immotile cells in bulge area (red) and monolayered regions (blue) plotted against time in active Brownian particle simulations. Solid lines and shaded areas denote the mean and standard deviation, respectively (N=16 simulation runs). The simulations started with all the particles being active, and 5% of the particles were set as passive starting at T = 30000 time steps.

## Discussion

In this study, we discovered that homogeneous *B. subtilis* swarms spontaneously develop a scattered pattern of interfacial liquid bulges, which consist of highly motile cells and maintain a constant difference (∼20% higher) in surface-packing cell density from elsewhere. The liquid bulges have a higher propensity to transit from motile to sessile biofilm state, presumably due to the enrichment of pre-existing immotile cells that naturally arise in a swarm. The formation of liquid bulges is correlated with local cell density fluctuation; it initiates when cell speed or local cell density exceeds a certain threshold, which is explained by an active Brownian dynamics model in terms of activity-induced pressure. Moreover, informed by the measured speed-density relation, we demonstrated reversible bulge formation by manipulating the speed and local density of cells with light. Our findings reveal a unique mechanism that promotes the motile-to-sessile state transition in surface-associated bacterial communities.

The liquid bulges we report here consist of almost equally motile cells and have a constant (∼20%) difference in surface-packing density of cells compared to elsewhere; these characteristics are distinct from those patterns driven by motility-induced phase separation (MIPS) (38) or by nucleation of immotile cells (39). We note that Grobas et al. reported that MIPS-induced clusters arising in the presence of stressors (e.g., the antibiotic kanamycin that collaterally reduces cell motility) could develop into ‘dynamic islands’ with highly motile cells packed in multiple layers (38). Despite the similarity between such dynamic islands and liquid bulges we report here, we stress that there are key differences between the two phenomena. First of all, liquid bulges in our phenomenon develop due to the increase of surface energy supported by activity-induced pressure rather than due to MIPS-induced clustering. Second, the further transition to chained biofilm cells is different in the two phenomena. In our case, the formation of chained biofilm cells in the bulges appears to be seeded by pre-existing sessile cells that are gradually enriched in liquid bulges; for MIPS-induced dynamic islands, it appears that the continuous increase of cell layer number or surface-packing density naturally gives rise to biofilms, whereas the liquid bulges in our case do not undergo multiple layering but maintain a constant difference in surface-packing cell density.

The colony mode of growth on solid surfaces is ubiquitous in environmental and clinical settings (77). Motile and sessile populations often coexist in bacterial colonies. In complex and fluctuating environments, the transition between the two states enables cells to optimize the growth of the entire population: Cells in sessile state will be more tolerant to unexpected environmental stresses (78, 79), while cells in motile state can explore more space and proliferate faster. Thus the motile-to-sessile transition promoted by bulge formation can be utilized as a bet-hedging strategy that enhances the adaptability of bacterial swarms and other surface-associated bacterial communities.

Our findings also suggest a new strategy, i.e., controlling liquid bulge formation, for biofilm pattern engineering. The ability to control biofilm patterns is essential for engineering bacterial living materials with desired functionalities (42-47). Informed by our study, we envisage that manipulation of cell motility and surface-packing density by either physical stimuli (e.g., light) or genetic control (80) may serve as a general means for biofilm patterning via liquid bulge formation. Moreover, our results show that the liquid bulges present a physical environment that tends to segregate passive particles from surroundings; the segregation is driven by unbalanced particle fluxes due to the differential diffusivity inside and outside the bulges. As such, manipulating liquid bulge formation enables phase separation or segregation in binary mixtures that consist of passive particles and an active matter bath, providing a novel approach for directed self-assembly of microscopic structures (48, 49).

## Materials and Methods

### Bacterial strains and preparation of swarm plates

The following strains were used: wild-type *B. subtilis* 3610; *B. subtilis* OMG991 with P_*hag*_-*gfp* reporter (*B. subtilis* 3610 *amyE*: : P_*hag*_-*gfpmut3* spec; gift from Harald Putzer (55)); *B. subtilis* DS91 (*sinI::spec* (60)); *B. subtilis* DK7921 (*sinI::kan amyE::*P*hag-gfpmut3 spec*; a *sinI*-deleted derivative of *B. subtilis* OMG991); *B. subtilis* 3610 with P_*tapA*_-*gfp* reporter (gift from Yunrong Chai (62)). To prepare bacterial swarms, single-colony isolates were grown overnight (∼10 to 12 h) with gyration at 180 rpm in LB broth (1% Bacto tryptone, 0.5% yeast extract, and 0.5% NaCl) at 30 °C. 10 mL swarm agar (0.6% or 0.8% Bacto-Difco agar infused with 1% Bacto tryptone, 0.5% Yeast Extract and 0.5% NaCl) was melted in a microwave oven, and poured to a polystyrene Petri dish (90 mm diameter, 15 mm height). The plates were placed inside a Plexiglas box with the lid off for 10 min.

After that, 1-2.5 μL diluted overnight culture (OD600 ∼ 1) were inoculated at 1.5 cm from the edge of the plate. Swarm plates were incubated at 30 °C and ∼50% relative humidity for 6-7 hours before imaging. For characterization of cell density, the membrane stain FM4-64 (Life Technologies, Cat. No. T13320) dissolved in deionized water was added at a final concentration of 1.0 μg/ml into the molten agar before it was poured into Petri dishes.

### Phase contrast, darkfield and fluorescence imaging

Imaging was performed on motorized inverted microscopes (Nikon TI-E or Nikon TI2-E). The following objectives were used in different experiments for phase contrast, darkfield, and fluorescence imaging: Nikon CFI Plan Fluor DL4×, N.A. 0.13, W.D. 16.4 mm; Nikon CFI Achromat DL 10×, N.A. 0.25, W.D. 7.0 mm (with a phase plate suitable for phase contrast imaging); Nikon CFI Plan Fluor 10×, N.A. 0.30, W.D. 16 mm (without phase plate and only for dark field imaging); Nikon CFI Super Plan Fluor ELWD ADM 20×C, N.A. 0.45, W.D. 8.2–6.9 mm; and Nikon CFI Super Plan Fluor ELWD ADM 40×C, N.A. 0.60, W.D. 3.6–2.8 mm. For darkfield imaging, the 4× objective was used to take the large-field image of the bulge pattern in Fig. 1 by stitching overlapping images; the 10× (N.A. 0.30, W.D. 16 mm) objective was used in other experiments. Fluorescence imaging was performed in epifluorescence using filter sets specified in sections below, with the excitation light provided by a mercury precentered fiber illuminator (Nikon Intensilight) or a solid state light engine (Lumencor SPECTRA X). Recordings were made with an sCMOS camera (Andor ZYLA 4.2 PLUS USB 3.0 or Andor Neo 5.5; Andor Technology) using the software NIS-Elements AR (Nikon). In all experiments the petri dishes were covered with lid to prevent evaporation and air convection, and the sample temperature was maintained at 30 °C using a custom-built temperature control system installed on microscope stage, unless otherwise stated.

### Single-cell motion pattern inside and outside liquid bulges

To measure the motion pattern of cells inside and outside liquid bulges, the swarm plates were prepared following the procedures as described above. Overnight culture of non-fluorescent cells (*B. subtilis* 3610 or *B. subtilis* DS91) was mixed with 0.5% fluorescent cells (*B. subtilis* OMG991 or *B. subtilis* DK7921) and the cell number density of the mixture was adjusted to OD600 ∼1 with fresh LB medium. The swarm plates were inoculated with 1-2.5 μL of the mixture. After incubation at 30 °C and ∼50% relative humidity for 6-7 hours, the swarm plates were then transferred to the microscope stage (Nikon TI-E). The motion of fluorescent cells was imaged in epifluorescence using the 20× objective and a Yellow Florescent Protein (YFP) single-band filter set (excitation 500/20 nm, emission 535/30 nm, dichroic: 515 nm; 49003-ET-EYFP, Chroma), with the excitation light provided by the mercury precentred fibre illuminator (Nikon Intensilight); this choice of excitation wavelength instead of regular GFP excitation wavelength at ∼480 nm is to reduce the photodamage to the cell. Meanwhile, the background bacterial motion in the swarm was imaged in phase contrast through the same optical system, with the illumination light provided by a white light LED (Cat. No. MCWHL5; Thorlabs). Recordings were made for a duration of 30 s by the Andor Zyla sCMOS camera. The camera was configured to record at 40 fps and exposure time for each frame was set as 15 ms; fluorescent and phase contrast images were recorded in alternate frames at 20 fps. The fluorescence excitation light was continuously illuminated and the white light LED was switched on only during the acquisition of phase contrast images. The camera was controlled by NIS Elements (Nikon); the white light LED was triggered by 20 Hz TTL signals sent from a custom-programmed Arduino microcontroller that modulated the 40 Hz fire output from the camera.

### Imaging of gene expression level during the development of liquid bulges

The swarm plates of *B. subtilis* 3610 strains with P_*hag*_-*gfp* reporter or P_*tapA*_-*gfp* reporter were prepared following the procedures described above. Both plates were mixed with 1.0 μg/mL FM4-64 to quantify the surface-packing density of cells. After the emergence of stable liquid bulges, we monitored the physiological state of cells inside and outside liquid bulges for ∼3 hr by imaging the three fluorescence reporters through the 10× phase contrast objective on a motorized microscope (Nikon TI2-E). FM4-64 is subject to rapid photobleaching under blue light illumination (i.e. excitation light of GFP), so it cannot be imaged simultaneously with GFP reporters in the same field of view. To overcome this difficulty while acquiring the cell-density corrected gene expression level for single cells, we first measured the fluorescence intensity distribution of GFP (from gene expression reporters) in a rectangular region (1.4 mm × 1.7 mm; Region 1) whose shorter side was parallel to the orientation of a stripe-like bulge; the selected area included monolayered regions outside the stripe. We then measured the fluorescence intensity distribution of FM4-64 in another rectangular region (Region 2) adjacent to and of the same dimensions as the one for GFP reporter imaging. The distributions of cell mass and of physiological state were nearly identical in these two adjacent regions, and thus the FM4-64 fluorescence distribution measured in one region can be used to compute the cell-density corrected gene expression level in the other region. Microscopy images were acquired for 160 min in 20 min intervals. At the beginning of each time interval, the following image acquisition procedures were performed automatically (controlled by software NIS-Elements AR; Nikon) and completed within ∼1 min: (1) Move the motorized stage to Region 1, perform autofocusing in phase-contrast channel, and acquire a phase-contrast image; (2) Acquire a fluorescence image of GFP reporter at Region 1; (3) Move the motorized stage to Region 2, perform autofocusing in phase-contrast channel, and acquire a phase-contrast image; (4) Acquire a fluorescence image of FM4-64 in Region 2. For the acquisition of fluorescence images, the excitation light was provided by a solid state light engine (Lumencor SPECTRA X), and the following filter sets were used: an enhanced green fluorescent protein (EGFP) filter set for P_*hag*_-*gfp* reporter and P_*tapA*_-*gfp* reporter [excitation 470/40 nm, emission 525/50 nm, dichroic: 495 nm; 49002-ET-EGFP (FITC/Cy2); Chroma]; and an mCherry filter set for FM4-64 (excitation 565/55 nm, emission 645/75 nm, dichroic: 600 nm; 49055-ET-Wide mCherry/Texas Red for 540-580nm LEDs; Chroma). For the acquisition of phase-contrast images, the transmitted light was provided by a built-in white light LED. All recordings were made by the Andor Neo sCMOS camera; the camera exposure time per frame for the fluorescent images of P_*hag*_-*gfp* reporter, P_*tapA*_-*gfp* reporter, and FM4-64 was set as 500 ms, 2 s and 1 s, respectively, while the exposure time per frame for phase-contrast images was set as 20 ms. The control of light sources and the camera were all performed by the software NIS-Elements AR (Nikon).

### Measurement of collective velocity in swarms

To measure the collective speed of cells during the development of liquid bulges (Fig. 2), phase contrast image sequences were acquired for 20 s and at 40 fps (with an exposure time of 15 ms for each frame) through the 4× phase contrast objective. The image sequences were taken every 20 minutes and for a duration of ∼3 hours. To measure the collective speed of cells during violet light illumination (Fig. 3A), dark field image sequence was acquired for 60 s and at 20 fps (with an exposure time of 20 ms for each frame) through the 10× dark field objective and a built-in 1.5× relay lens of the microscope. All recordings were made by an sCMOS camera (Andor Zyla) and controlled by the software NIS-Elements AR (Nikon). The collective speed was obtained by optical flow analysis on the image sequences (see section “Image processing and data analysis” below).

### Violet light control of liquid bulges

To study the effect of cell motility and local cell density variation on liquid bulges (Fig. 3A), a circular area of ∼7 mm^2^ in a swarming colony was illuminated by violet light at an intensity of ∼6 mW/mm^2^ for ∼25 seconds. To trigger bulge formation by increasing surface-packing density of cells in a swarming colony (Fig. S6), the colony was illuminated by violet light at an intensity of ∼2-3 mW/mm^2^ for 1-2 min. In these experiments, violet light illumination was provided by the mercury precentred fibre illuminator (Nikon Intensilight) through a single band filter set (excitation 406/15 nm, emission 535/22 nm, dichroic: 458 nm; Semrock Inc.) and the 10× darkfield objective.

### Single-cell motion pattern during violet light illumination

To investigate the motion pattern of individual cells during violet light illumination, we collected *B. subtilis* 3610 swarm cells by depositing 20 μL LB broth to the edge of a swarming colony and transferring the ∼20 μL cell suspension to an Eppendorf tube. The collected cell suspension was diluted by 50-fold with LB broth. Then the diluted suspension was deposited to fresh agar surface as a ∼1 mm diameter liquid drop containing hundreds of cells following bubble collapse. The motion of cells in the suspension drop was recorded for 60 s in phase contrast by an sCMOS camera (Andor Zyla) through the 10× phase contrast objective. The camera was configured to record at 20 fps and the exposure time for each frame was set as 20 ms. Violet light illumination was provided by the mercury precentred fibre illuminator (Nikon Intensilight) through a single band filter set (excitation 406/15 nm, emission 535/22 nm, dichroic: 458 nm; Semrock Inc.). When switched on, violet light was illuminated at an intensity of ∼4 mW/mm^2^ to the entire area accessible to cells confined in the suspension drop.

### Relation between cell motility and surface-packing density

To measure the speed-density relation, overnight cultures of *B. subtilis* DS91 and fluorescent non-chaining mutant *B. subtilis* DK7921 (Δ*sinI* P_*hag*_-*gfp*) were adjusted to OD600 ∼1 with fresh LB medium. Then 5% *B. subtilis* DK7921 cells were mixed with *B. subtilis* DS91 cells. Swarm plates were inoculated with this mixture and incubated following the procedures described above, before being transferred to the microscope stage (Nikon TI-E). The optical setup, as shown in Fig. S5, integrates large-scale (∼3 cm^2^) blue/violet light illumination (for speed reduction in the swarm), darkfield or phase contrast microscopy (for speed measurement by optical flow analysis), and wide-field epifluorescence microscopy (for surface-packing density measurement by counting the number of fluorescently labeled cells). Violet light (395/25 nm) and blue light (440/20 nm) are used for reducing the cell speed, while green light (550/15 nm) is the light source of dark field or phase contrast microscopy. All of the three wavelengths are from a solid light engine (Lumencor SPECTRA X) and the light beams pass through a custom-built phase ring and a condenser from above the stage. The custom-built phase ring is a key component that enables the large-field speed control and dark field microscopy. The phase ring consists of a shortpass filter (500 nm; diameter ∼2.5 cm; Kaitao OpticSc Compony Limited) placed at the center and on top of a flat glass lens (diameter ∼3.3 cm; from Kaitao OpticSc Compony Limited); both the shortpass filter and the flat glass lens are installed in a 3D-printed holder (outer diameter ∼3.6 cm, inner diameter ∼3.3 cm; made from polylactide (PLA)). The shortpass filter only allows the blue and violet light in the setup to pass through, and thus the green light can only pass through the remaining annulus region of the flat glass lens. Below the stage, light from the mercury illuminator (Nikon Intensilight) transmits through an FITC filter set (excitation 482/35 nm, emission 536/40 nm, dichroic: 506 nm; FITC-3540C-000, Semrock Inc.) and serves as the excitation light for wide-field epifluorescence microscopy.

Microscopy images were acquired for ∼10-20 min in 1 min intervals. At the beginning of each time interval, the following image acquisition procedures were performed and completed within ∼40 s (including the time taken for data transfer from camera’s memory buffer to computer storage): (1) Take a 5-second dark field image sequence (80 fps, 6 ms exposure time for each frame) at a selected region of interest (ROI) (1024 pixel × 1024 pixel, or 660 μm × 660 μm) for the measurement of collective cell speed by optical flow analysis (see below); (2) Take one fluorescence image for cell number counting (100 ms exposure time for each frame); (3) Take one dark field image over the entire field of view for manual identification of the bulge area (20 ms exposure time for each frame). Recordings were made by an sCMOS camera (Andor Zyla) through the 10× dark field objective. The violet and blue light was kept on in the whole recording process. The green light and the fluorescence excitation light were only switched on during the acquisition of darkfield and fluorescence images, respectively. The green light for dark field imaging was triggered by a TTL signal sent from a custom-programmed Arduino microcontroller, while the excitation light from the mercury illuminator below the stage was controlled by a custom-built mini-robotic arm [consisting of a servo motor (SG90; Tower Pro) and a 3D-printed handle made of PLA] controlled by pulse-width modulation (PWM) signal sent from another custom-programmed Arduino microcontroller; both Arduino microcontrollers received signals from a custom-written macro in NIS-Elements AR (Nikon) that controlled the entire image acquisition protocol.

We note that the fraction of fluorescent cells in swarming colonies may differ from that at the initial inoculum and vary in space due to slight growth rate difference. To convert the surface-packing density of fluorescent cells to the absolute cell surface-packing density, another batch of microscopy images was acquired. These microscopy images were acquired for ∼10-20 min in 1 min intervals, and recordings were made by an sCMOS camera (Andor Zyla) through the 40× objective for better cell segmentation. At the beginning of each time interval, the following image acquisition procedures were performed and completed within ∼20 s (including the time taken for data transfer from camera’s memory buffer to computer storage): (1) Take a 5-second phase contrast image sequence (80 fps, 6 ms exposure time for each frame) at a selected region of interest (ROI) (1024 pixel × 1024 pixel, or ∼166 μm × ∼166 μm) for the measurement of collective cell speed by optical flow analysis (see below); (2) Take one phase contrast image over the entire field of view for manual identification of the bulge area and for counting the absolute number of cells (20 ms exposure time for each frame). In the whole imaging process, the violet and blue light was kept on. The green light was only switched on during the acquisition of phase contrast images and triggered by a TTL signal sent from a custom-programmed Arduino microcontroller, which received signals from a custom-written macro in NIS-Elements AR (Nikon) that controlled the entire image acquisition protocol.

### Microsphere dynamics during bulge formation

To measure the motion and distribution of passive microspheres, we diluted a microsphere suspension (Polybead Sulfate Microspheres, 1 μm; Polysciences, Cat. No. 19404-15) by 1000-fold in the LB broth to a final concentration of 4.55×10^7^ particles/mL, and deposited the suspension to a swarming colony by bubble collapsing. The deposited microspheres underwent diffusive motion in the swarm due to advection by motile cells. They were allowed to disperse from the initial drop position for ∼30 min until reaching a relatively homogeneous density distribution in the entire field of view. When liquid bulges emerged in the field of view, the microspheres and the colony were recorded continuously for 8 min in dark field microscopy through the dark field 10× objective, and recordings were made by the Andor Zyla sCMOS camera at 20 fps (20 ms exposure time for each frame).

### Image processing and data analysis

Images were processed using the open-source Fiji (ImageJ) software (http://fiji.sc/Fiji) and custom-written programs in MATLAB (The MathWorks; Natick, Massachusetts, United States).

To identify the boundary and area of liquid bulges, we made use of the fact that, due to frequent cell-cell overlapping, the liquid bulges have a greater brightness in dark field images or have a different contrast pattern in phase contrast images compared to that in the monolayered regions. For Fig.1, 2 and 3B and Movie S2, the boundary of liquid bulges was determined manually. In the study of the effect of cell motility and local cell density variation on liquid bulges (Fig. 3A), we detected and computed the area of liquid bulges in a darkfield image by thresholding and binarization, with the threshold being a constant plus the mean grayscale value of the image (a constant threshold was not applicable to all images due to the illumination of violet light). In the study with passive microspheres (Fig. 5), we identified the area of bulges in a darkfield image following the steps below: (1) Reduce non-uniformity of light illumination and segment bulges by a bandpass filter with a custom-written program in MATLAB (based on the algorithm of bandpass filter in ImageJ); (2) Smooth the image with Gaussian filter; (3) Threshold and binarize the image with an appropriate threshold.

For single-cell or single-microsphere motion pattern analysis, the trajectories of cells or microspheres were obtained by a particle-tracking program in MATLAB described earlier (81). Prior to tracking, the images were processed and binarized following the procedures described above for identifying the area of bulges in the presence of passive microspheres. To improve the accuracy of motion pattern analysis, we removed short trajectories (<2.5 s) that could result from tracking errors. The instantaneous velocity of a cell or a microsphere was calculated from the displacement of the center of the cell/microsphere between two consecutive frames. The mean speed of cells was then obtained by taking the temporal and ensemble average of cell speed during the entire tracking period. The mean speed of microspheres (Fig. 5B) at time *t*_0_, denoted as 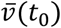, was computed by averaging all the instantaneous speeds of all microspheres at *t*_0_ over a 1-s time window [*t*_0_-0.5 s, *t*_0_+0.5 s]. The probability density function (PDF) of speed distributions was derived from the computed instantaneous speeds. The velocity autocorrelation function (VACF) is defined as 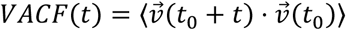, where 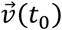 is the instantaneous velocity of a cell/microsphere at time *t*_0_ and the angular brackets denotes the ensemble average over all cells/ microspheres. To compute the normalized VACF, *VACF*(*t*) was normalized by *VACF*(0). For a particle undergoing diffusive motion, VACF follows an exponential decay, i.e., 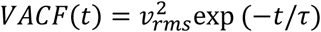, where *τ* is the persistence time and *ν*_*rms*_ is the root-mean-square speed; the diffusivity *D* can be calculated as 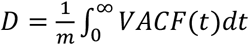, where m is the dimension (for a quasi-2D motion in a bacterial swarm, m = 2). Thus we obtained *τ* and *ν*_*rms*_ by least-squares fitting of the VACF, and obtained the diffusivity by integrating the VACF. Mean square displacement (MSD) was calculated as *MSD*(*t*) = ⟨|*x*(*t*_0_ *+ t*) *−x*(*t*_0_)|^*2*^⟩, where *x*(*t*) is the position of a cell at time *t* and the angular brackets denotes ensemble average over all cells. The tumble rate of cells (Fig. S3) at a specific time is defined as the number fraction of cells that are undergoing tumbling events; here a tumbling event is defined by the rate of change of velocity direction greater than 30°/frame or by an angular speed >∼10.5 rad/s (82). Detection of tumbling events based on speed decrease is less sensitive than that based on the angular speed for *B. subtilis* swarm cells, because cells only slightly decreased speed during re-orientation.

For the calculation of collective cell speed (Fig. 2A and Fig. 3A, B), we performed optical flow analysis based on phase contrast or darkfield microscopy images using the built-in function opticalFlowFarneback of MATLAB. The optical flow field for any two consecutive video frames was estimated using Farneback method (83). The grid size of the optical flow field was set as 1 pixel × 1 pixel. The obtained optical flow fields were used to compute the collective cell speed in Fig. 2A and Fig. 3A, B. For the optical flow fields used in Fig. 2, the parameters of optical flow calculation were set as follows: number of pyramid layers, 3; image scale, 0.5; number of search iterations per pyramid level, 5; size of the pixel neighborhood, 5; averaging filter size, 4. For the optical flow fields used in Fig. 3A,B, the parameters of optical flow calculation were set as follows: number of pyramid layers, 3; image scale, 0.5; number of search iterations per pyramid level, 3; size of the pixel neighborhood, 5; averaging filter size, 15.

To compute the single-cell gene expression level during the development of liquid bulges based on the fluorescence images of P_*hag*_-*gfp* reporter, P_*tapA*_-*gfp* reporter, or FM4-64, we selected from a fluorescence image two regions of interest (ROI) (∼100 - 300 μm in width, ∼450 - 700 μm in length) whose longer sides were parallel to the orientation of the stripe-like bulge; one ROI was selected from monolayered regions and the other from the bulge. All subsequent image analysis was performed within the ROIs. To obtain a reporter or FM4-64 fluorescence image *I*_*c*_(*t*) at time *t* corrected for the fluorescence background and illumination inhomogeneity of excitation light, we used the formula *I*_*c*_(*t*) = [*I*_0_(*t*) *−I*_*b*_(*t*)]/*I*_*e*_(*t*), where *I*_0_(*t*), *I*_*b*_(*t*), and *I*_*e*_(*t*) are the gray-scale image at time *t* of the fluorescent reporters, of the associated background fluorescence from the agar substrate, and of the normalized background fluorescence, respectively. To obtain the background fluorescence image *I*_*b*_(*t*) associated with *I*_0_(*t*), we first acquired a time series of fluorescence images 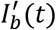 on cell-free agar surfaces following the same imaging protocol described in section “Imaging of gene expression level during the development of liquid bulges”. For P_*hag*_-*gfp* and FM4-64 fluorescence images, to account for the variability of background fluorescence across different experiments while acquiring *I*_0_(*t*), we computed *I*_*b*_(*t*) as 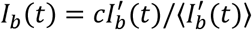, where 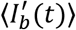 is the spatial average of pixel intensities of 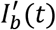 and *c* is a scalar constant that varied for different time series of *I*_0_(*t*) but remained unchanged throughout the same time series, which ensures that the minimal pixel value of (*I*_0_ *−I*_*b*_) throughout the entire time series is close to 0 (i.e., this minimal pixel value is regarded as the fluorescence intensity from cell-free background). The normalized background fluorescence image *I*_*e*_(*t*), was computed as 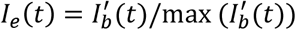, where max 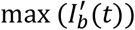 denotes the maximal pixel value of 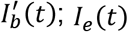; *I*_*e*_(*t*) served to correct for the illumination inhomogeneity of excitation light. For the time series of P_*tapA*_-*gfp* fluorescence images *I*_0_(*t*), we noted that the pixel values of (*I*_0_ *−I*_*b*_) at *t*= 0 min as computed by the procedures above was almost zero throughout the entire image, suggesting that the expression level of TapA was negligible at the initial state of bulge development. Making use of this fact, we chose to use *I*_0_(*t* = 0) as a genuine fluorescence background for more accurate correction of all P_*tapA*_-*gfp* fluorescence images in a specific time series: first, to account for the photo-bleaching of background fluorescence within the time series *I*_0_(*t*), we computed *I*_*b*_(*t*) as *I*_*b*_(*t*) = *γ*(*t*)*I*_0_(*t* = 0)/*γ*(*t* = 0), where *γ*(*t*) is a time-dependent scalar parameter defined as the spatially averaged pixel value of a uniform monolayered region within the image *I*_0_(*t*); next, to correct for the illumination inhomogeneity of excitation light, the normalized background fluorescence image *I*_*e*_(*t*) was computed as *I*_*e*_(*t*) = *I*_0_(*t* = 0)/max (*I*_0_(*t* = 0)). Having obtained the background- and illumination inhomogeneity-corrected reporter or FM4-64 fluorescence images *I*_*c*_(*t*) in the two ROIs, the single-cell gene expression level inside and outside liquid bulges was computed as ⟨*I*_*c*_(*t*)⟩_reporter_/⟨*I*_*c*_(*t*)⟩_FM4−64_, where angular brackets indicate spatial average of pixel values over the entire image. Finally, the spatial variation of reporter activity at time *t* was computed as the standard deviation of the pixel value distribution in *I*_*c*_(*t*) divided by the spatially average pixel value of *I*_*c*_(*t*), which is equivalent to the coefficient of variance of the pixel value distribution. The image processing was done with a custom-written program in MATLAB.

To compute the cell density fluctuation *ΔN/N* in monolayered regions in Fig. 3A, we assumed that cells have uniform size and thus used the area occupied by cells as a convenient measure of cell number. We first computed the cell area in each darkfield microscopy image by thresholding and binarizing the grayscale image, with the threshold being a constant plus the mean grayscale value of the image. Then we divided each image into a series of non-overlapping interrogation windows of a size comparable to the transient bulge-like domains with high surface-packing cell density (32 pixel × 32 pixel, or ∼14 μm^2^). For the i-th interrogation window, we obtained a time series of the cell area *n*_*i*_(*t*) over all image frames. The cell density fluctuation at a specific time *t* was then calculated as the standard deviation (*ΔN*) of {*n*_*i*_(*t*), *i* = 1,*2*, … } divided by the mean *N*; the resultant instantaneous cell density fluctuation was further averaged over a 1-s time window comparable to the duration of transient domains.

To compute the surface-packing density of cells when measuring the speed-density relation of cells (Fig. 3B,C), we first counted the number of fluorescently labeled cells in fluorescence images by the built-in ImageJ functions *background subtraction* (based on rolling ball algorithm) and *Find Maxima* (based on H-maxima transform (84)). The fluorescent cell number density inside or outside liquid bulges was calculated as the fluorescent cell count divided by the corresponding area. To convert the surface-packing density of fluorescent cells to the absolute cell surface-packing density, we acquired a separate batch of phase contrast microscopy images through the 40× objective for direct counting of all cells (see section “Relation between cell motility and surface-packing density” above). Based on these high-resolution phase contrast images we calculated the absolute number of cells in monolayered regions (i.e., outside liquid bulges) as the total area occupied by cells divided by the single-cell area; here the total cell area was computed by binarizing the image with an appropriate threshold, and the single-cell area was estimated by taking the average of those segmented cell areas of size in the range of ∼3-8 μm^2^. The absolute surface-packing density of cells in monolayered regions with intermediate collective cell speeds (∼30-40 μm/s) was then used to calibrate the surface-packing density of fluorescent cells in monolayered regions with the same range of collective cell speeds, yielding the true ratio between fluorescent cells and non-labeled cells at the region of interest that differed from the initial mixing ratio due to growth rate difference. The absolute surface-packing cell density in Fig. 3D was estimated following the same procedures described above, except that darkfield images acquired through the darkfield 10× objective and the built-in 1.5× relay lens of the microscope were used.

### Active Brownian dynamics model

We modeled cells as self-propelled rod-like particles moving in a quasi-2D square-like area of size *L* × *L* and with periodic boundary conditions. Each modeled cell consists of n circular disks of diameter *d* whose centers are separated by a fixed distance *Δ* (85). The disk diameter and inter-center separation were chosen to make the aspect ratio of a rod ∼5:1, roughly corresponding to the aspect ratio of *B. subtilis* cells. The center position ***r***_***i***_ and orientation *θ*_*i*_ of the *i*-th cell follow overdamped Langevin dynamics (73):

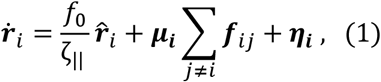

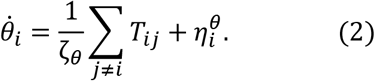

Eqn. (1) and (2) describe translational and rotational motion of the cell, respectively. Here *f*_0_ is the self-propulsive force due to the hydrodynamic interaction between rotating flagella and the swarm fluid film; 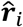 is a unit vector with an orientation angle *θ*_*i*_; 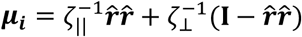is the mobility tensor, with *ζ*_||_ and *ζ*_*⊥*_ being the translational drag coefficient in the direction parallel and perpendicular to the cell orientation, respectively; *ζ*_*θ*_ is the rotational drag coefficient given by *ζ*_*θ*_ = (1 *+ a*^*2*^) × *ζ*_*⊥*_/1*2* (*a* is aspect ratio) (86). ***η***_***i***_ is the translational noise given by ***η***_***i***_ = ***μ***_***i***_ ***f***_***n***_, with ***f***_***n***_ being a random force vector whose x- and y-components are independent and follow the same Gaussian distribution 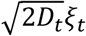; here *D*_*t*_ and *ξ*_*t*_ are translational diffusion constant and a Gaussian white noise with zero mean and unit variance, respectively. 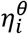 is the rotational noise given by 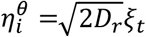, with *D*_*r*_ and *ξ*_*r*_ being the rotational diffusion constant and a Gaussian white noise with zero mean and unit variance, respectively. The interaction force ***f***_*ij*_ is modeled as harmonic repulsion given by 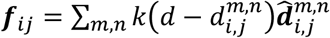 for 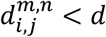 and ***f***_*ij*_ = 0 otherwise, where *k* is the strength coefficient of steric interaction between cells,*d* is the interaction range, 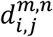and 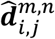 are the distance and the unit vector from the *n*-th disk of cell *j* to the *m*-th disk of cell *i*, respectively. The torque *T*_*ij*_ is given by 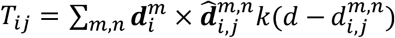, where 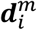 is the vector from the center position of cell *i* to the *m*-th disk of the cell.

Using the model we first established the relation between activity-induced pressure and cell’s active speed 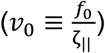 as well as the surface-packing density of cells (*ρ*_0_) in the simulation of cell monolayers. Flagellated bacteria can be regarded as force dipoles exerting active stresses to the surrounding fluid (87); for cells in the swarm fluid film, the active stress component acting on the air-fluid interface must be proportional to that acting laterally in the plane of the fluid film. Therefore we considered cells confined by a square domain (of size ∼1600 cell area, or 80 × 80 unit area) with hard confinement walls and took the activity-induced pressure exerted to the lateral confinement walls as a proxy for the activity-induced pressure exerted on the air-liquid interface. The interaction between cells and the confinement walls was modeled as harmonic repulsion as defined above for cell-cell interactions. The activity-induced pressure was defined as the average force exerted on the lateral confinement walls per unit length. By scanning over 2040 parameter sets of (*ν*_0_,*ρ*_0_), we obtained the contour lines of activity-induced pressure on the plane of *ν*_0_ and *ρ*_0_ after smoothing out the fluctuations.

To model the formation of liquid bulges, we introduced two additional variables in the model, local density *ρ*(*x, y*; *t*) and excess height *h*(*x, y*; *t*). At time *t*, the local density *ρ*(*x, y*; *t*) at position (*x, y*) is computed by averaging the cell density within a square region of side length *l* centered at (*x, y*), and the excess height *h*(*x, y*) in the range of (0,1] is used to identify whether a bulge can be formed in a certain area. When *ρ*(*x, y*; *t*) reaches a critical value *ρ*_*c*_ (i.e., the threshold of local surface-packing cell density for liquid bulge formation at a specific *ν*_0_ under the prescribed bulge-forming threshold of activity-induced pressure *π*_0_), *h*(*x, y*; *t*) is taken to be 1. When *ρ*(*x, y*; *t*) < *ρ*_*c*_, the activity-induced pressure becomes smaller than *π*_0_ and would not be able to support the bulging of the liquid film, and thus we let *h*(*x, y*; *t*) follow an exponential decay. To summarize, the dynamics of *h*(*x, y*; *t*) is described by the following function,

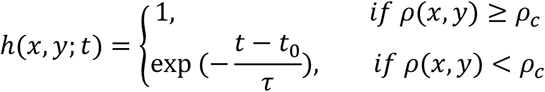

where *τ* is the relaxation time and *t*_0_ is the most recent time when *ρ*(*x, y*) became lower than *ρ*_*c*_. The bulge area is defined as the set of all positions where *h* is higher than 1/*e*. In experiments, cells’ motion in the liquid bulges is not restricted to two dimensions but can access the third dimension, and cells are able to overlap with each other. Thus cells in liquid bulges experience weaker steric interaction between each other. To model this fact, we set the strength of steric interaction *k* to be different inside and outside bulges as follows,,

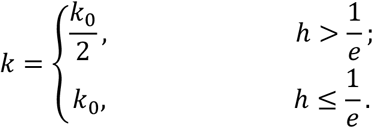

In addition, we observed in experiments that cells inside liquid bulges have a ∼7% smaller speed than elsewhere, presumably due to the differences in mechanical environment inside and outside bulges. To account for the observed speed difference in simulations, we set the effective self-propulsive force outside bulges to be 1.2-fold of that inside bulges, which yields a ∼5-10% speed difference of cells consistent with experimental measurements (Fig. S7).

For the simulation of bulge formation, once *π*_0_ is chosen, *ρ*_*c*_ at active speed *ν*_0_ can be determined from the contour plot (Fig. 4A). The active speed *ν*_0_ ranges from 1.0 to 5.0 unit length per unit time (with 1 unit time equal to 100 time steps), corresponding to a real speed of 30-150μm/s (i.e., 1 unit length and 1 unit time in simulations correspond to 1μm and 1/30 s in experiments, respectively). As another validation of the model, we found that the persistent time of modeled cells in simulations is ∼10 unit time, which is consistent with the persistence time ∼0.30 s measured in experiments. All analysis of simulation data was performed for the steady state after 40000 time steps (or 400 unit time). All parameters used in the simulations are summarized in Table 1.

**Table 1.**
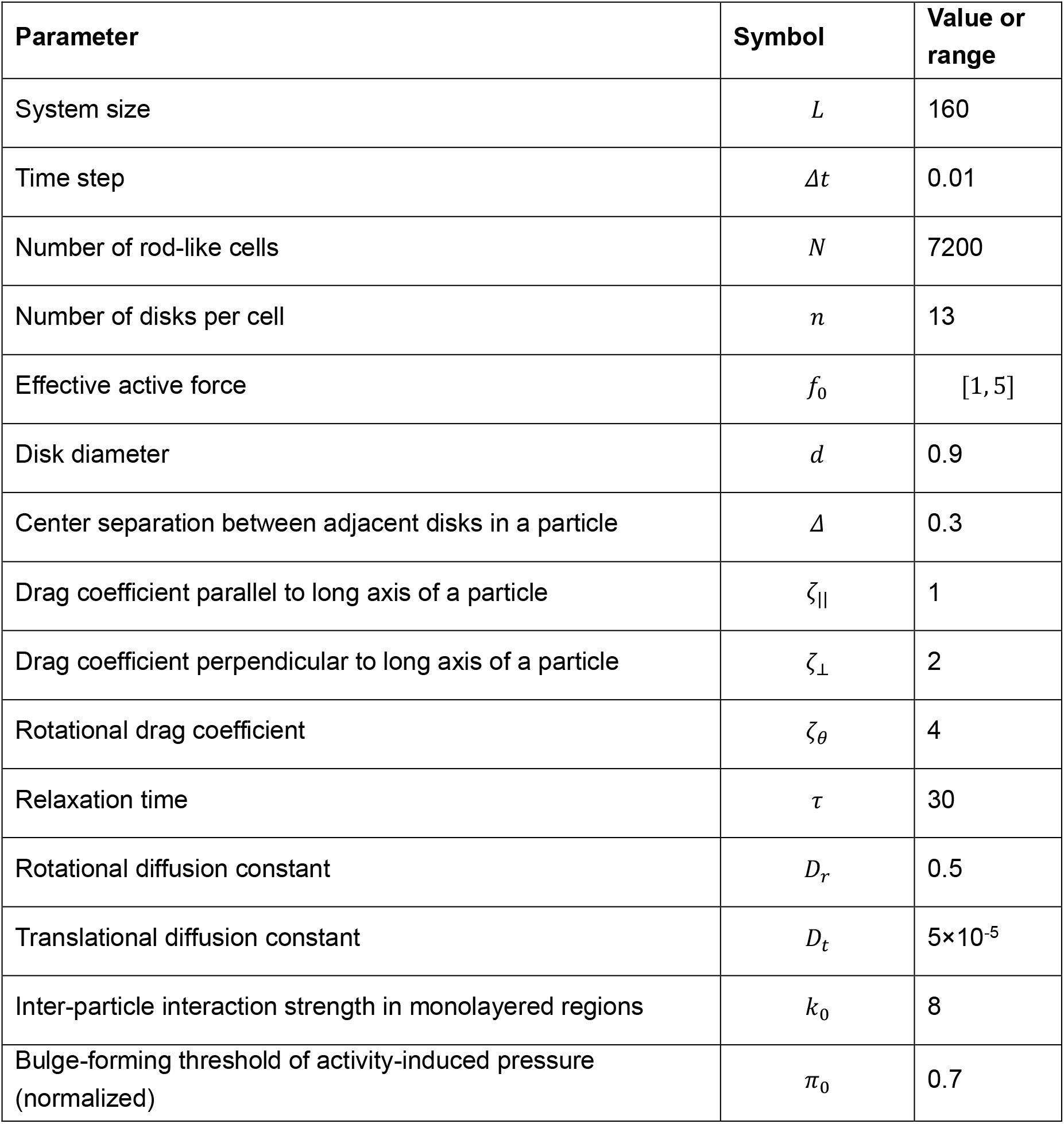
Simulation parameters.

For the simulation of enrichment of immotile cells in bulges, the simulations started with all the particles being active. 5% of the particles were then set to be passive (i.e., let *ν*_0_ = 0) starting at t= 30000 time steps, at which time bulges had already emerged. The simulation parameters used are the same as above.

## Acknowledgements

We thank Harald Putzer and Yunrong Chai for their kind gifts of bacterial strains; Wenlong Zuo for providing the custom-written single cell tracking program; Lei Xu for assistance with dark-field imaging; and Munehiro Asally and Marco Polin for helpful discussions. **Funding:** National Natural Science Foundation of China (NSFC No.

31971182, to Y.W.); Research Grants Council of Hong Kong SAR (RGC Ref. No. 14306820, RFS2021-4S04 and CUHK Direct Grants; to Y.W.);

National Institutes of Health of United States (R35 GM131783 to D.B.K).

## Data and materials availability

All data are available in the main text or the supplementary materials.

## Supplementary Materials for

**Other Supplementary Materials for this manuscript include the following:**

Movies S1 to S9

SM Figures

**Fig. S1.**
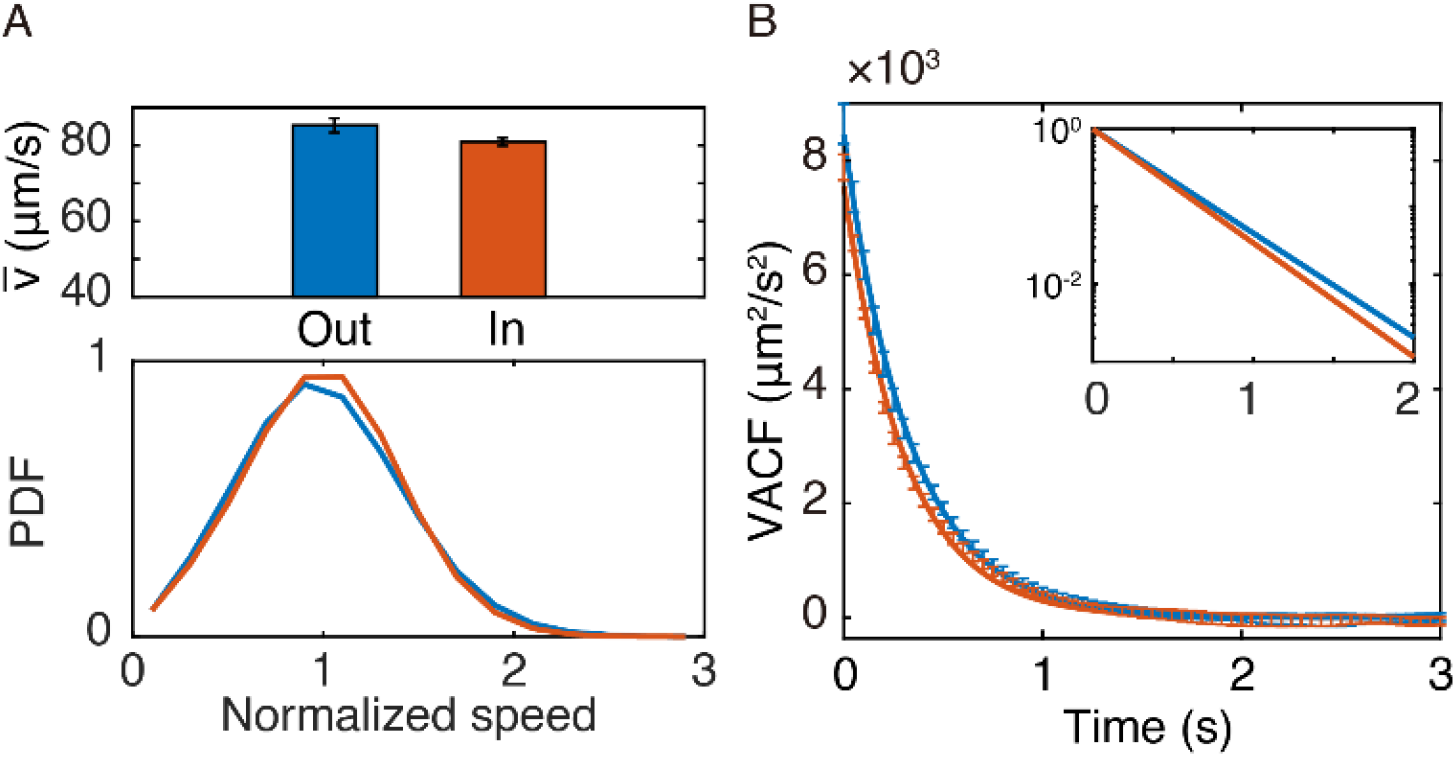
Motion pattern of B. subtilis ΔsinI cells inside and outside liquid bulges. (A) Speed of non-chaining *B. subtilis* Δ*sinI* cells inside and outside the liquid bulges. Upper panel: Average speed 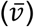 of cells that remained outside and inside the liquid bulges during the tracking period are shown as blue (85.3 ± 1.9 μm/s; mean ± SE, N = 12) and red (81.0 ± 1.0 μm/s; mean ± SE, N = 9) columns, respectively. Lower panel: Average probability density functions (PDFs) of the normalized speed of cells corresponding to the two scenarios in the upper panel, following the same color coding as that of the columns in upper panel. The speed was normalized by the mean speed of all cells in the corresponding scenario. (B) Velocity autocorrelation functions (VACFs) of cells that remained inside and outside the liquid bulges computed based on single cell trajectories. Solid lines are exponential fits to the VACFs [blue: outside liquid bulges, VACF_out_ ∼ exp(-t/0.33), persistence time *τ*_out_ = 0.33 ± 0.01 s; red: inside liquid bulges, VACF_in_ ∼ exp(-t/0.31), persistence time *τ*_in_ = 0.31 ± 0.01 s). Plotted in the inset are normalized fitted VACFs outside (blue) and inside (red) liquid bulges in semi-logarithmic scale. The diffusivity computed from VACF is ∼16% lower inside bulges than outside (∼1170 μm^2^/s vs. ∼1400 μm^2^/s) (Methods). Error bars represent the standard deviation of N=12 (blue) or 9 (red) biological replicates.

**Fig. S2.**
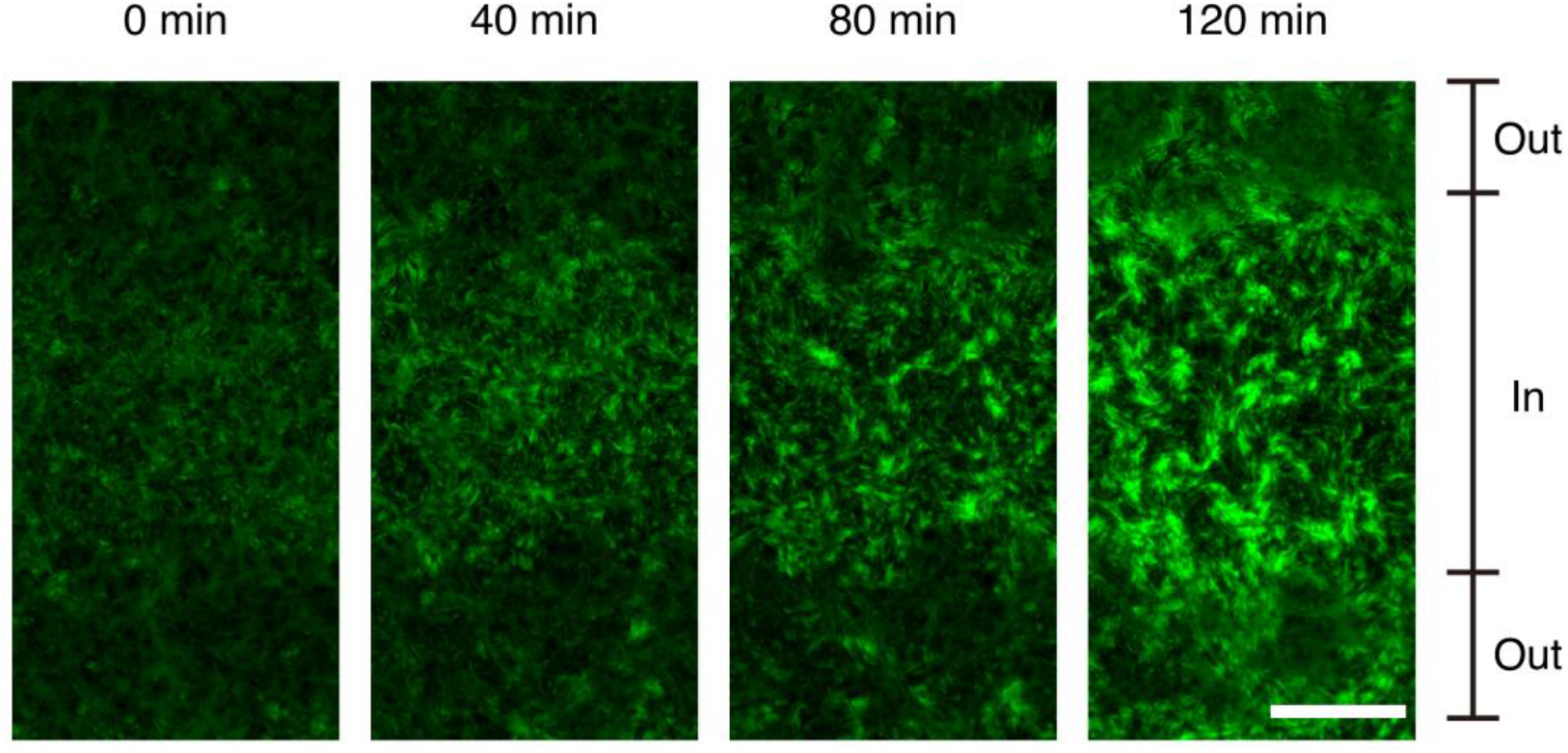
Spatial variation in Phag-gfp reporter activity. This fluorescence microscopy image sequence shows the development of P_*hag*_-*gfp* reporter fluorescence in pseudo-color inside and outside liquid bulges (as indicated by the labels on the right). Randomly spaced cellular clusters with higher P_*hag*_-*gfp* reporter fluorescence than elsewhere emerged inside bulges at the later stage of the development of stripe-like bulges. Scale bar, 200 μm.

**Fig. S3.**
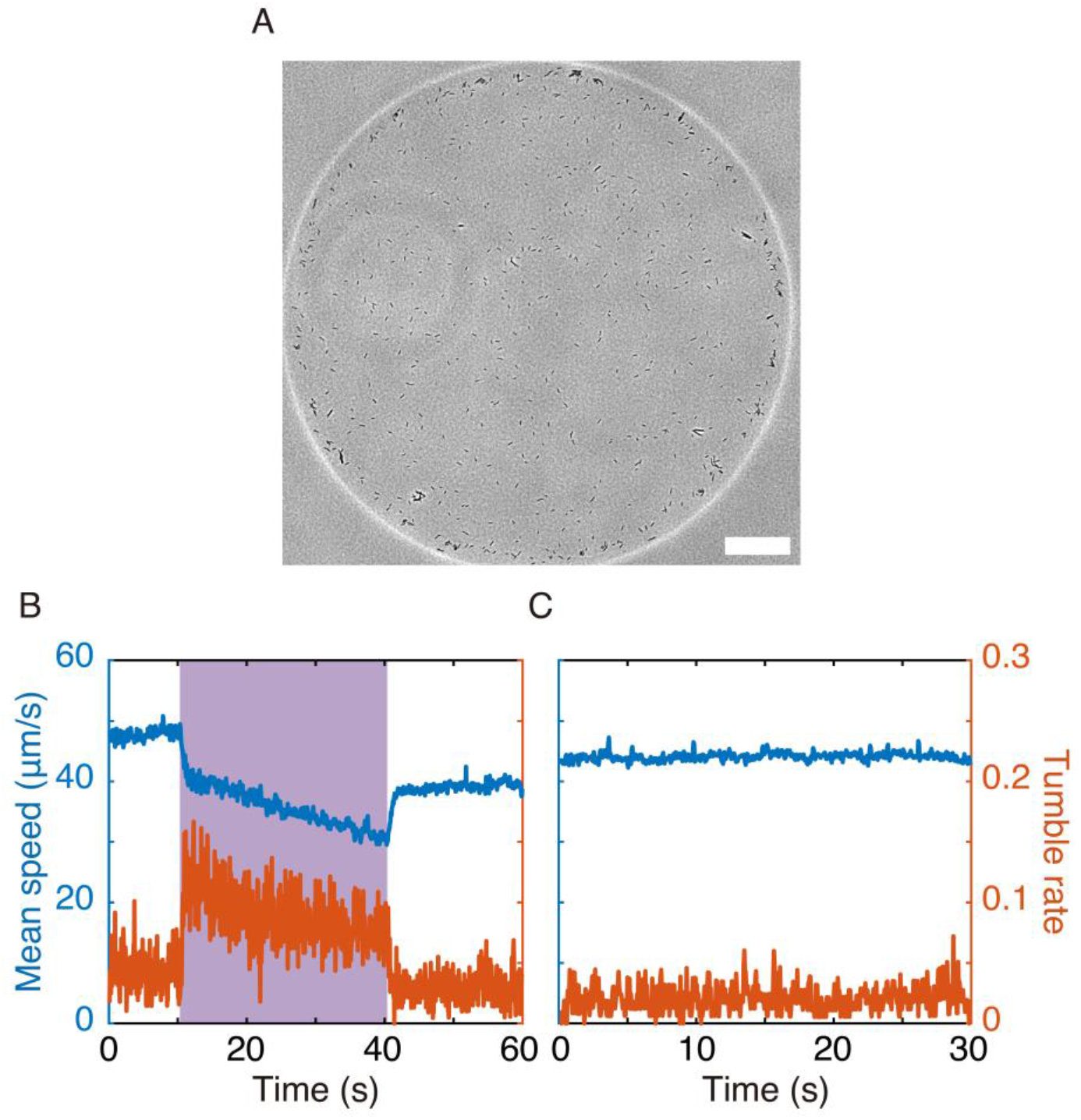
Reversible control of cell speed with violet light. (A) A representative phase-contrast microscopy image showing a quasi-two-dimensional suspension drop of low cell density for monitoring motion pattern of individual cells during violet light illumination (Methods). Scale bar, 100 μm. (B,C) Temporal dynamics of mean cell speed (blue) and tumble rate (red; Method) with (panel B) and without (panel C; as a control) violet light illumination (Methods). Violet light was illuminated during the period shaded in purple color in panel B. The results show that, as the violet light illumination was turned on, the speed of *B. subtilis* cells first dropped abruptly and then decreased slowly; upon retraction of violet light, the speed suddenly increased and remained constant at a level slightly lower than the original speed (i.e., not fully recovered to normal likely due to the permanent damage on cells by the violet light). Meanwhile, the tumble rate of cells experienced an abrupt increase when violet light illumination was turned on, followed by a slow decrease until returning to normal upon retraction of violet light illumination.

**Fig. S4.**
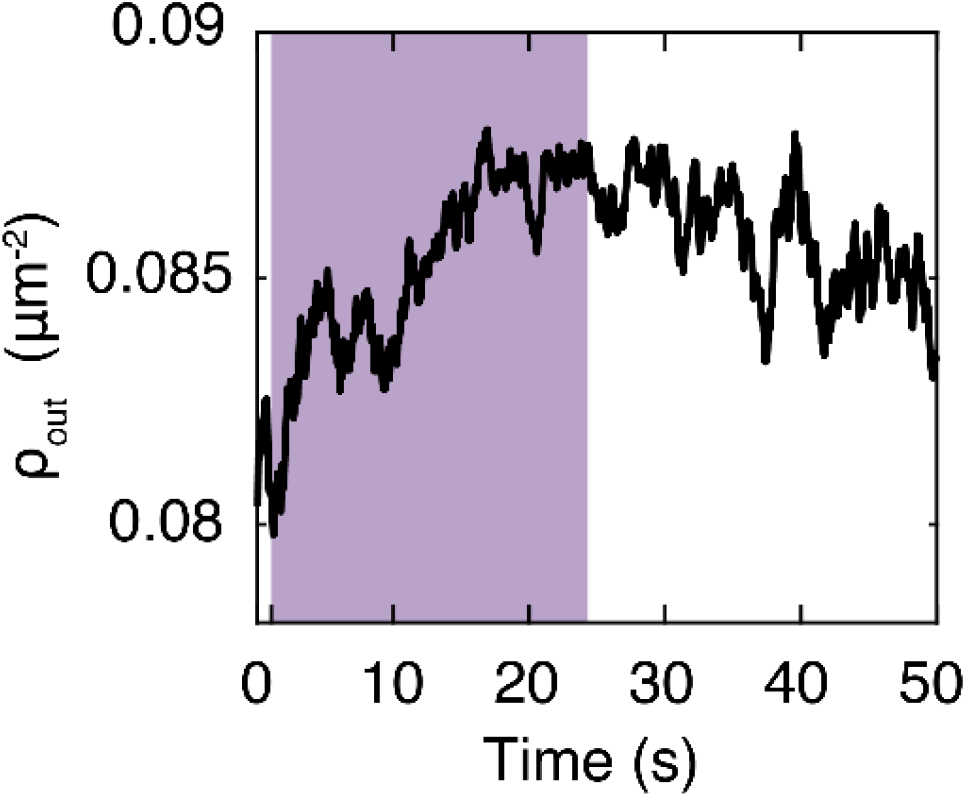
Temporal dynamics of surface-packing density of cells in a monolayered area during violet light illumination. Violet light was illuminated during the period shaded in purple color. The surface-packing density (*ρ*_*out*_) in the monolayered regions was increased by ∼10% at the end of violet light illumination.

**Figure S5.**
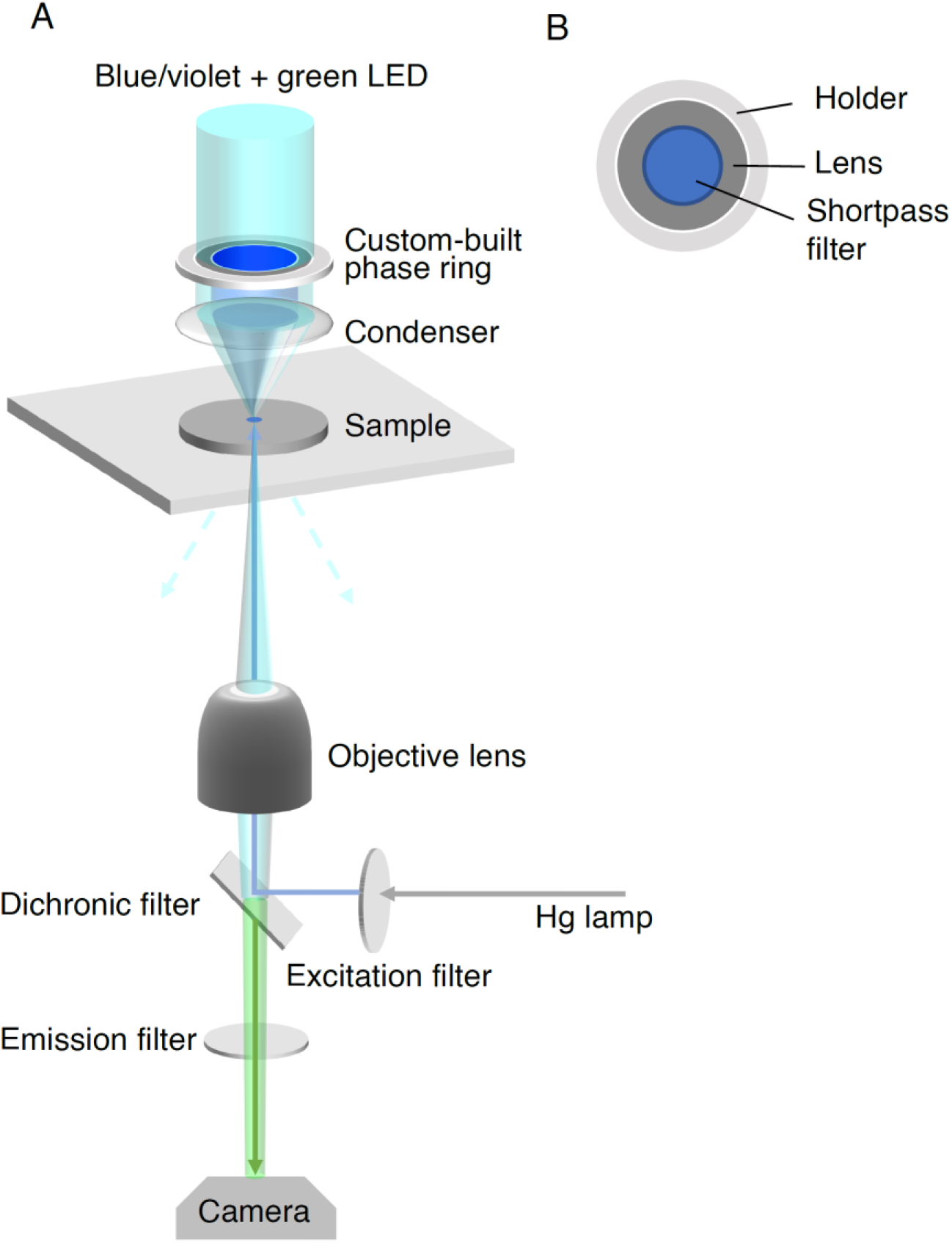
Schematic diagram of the integrated multi-mode imaging system for speed-density relation measurement. (A) The multi-mode imaging system integrates large-scale (∼3 cm_2_) blue/violet light illumination (for speed reduction in the swarm), darkfield microscopy (for speed measurement by optical flow analysis), and wide-field fluorescence microscopy (for surface-packing density measurement by counting the number of fluorescently labeled cells). The blue/violet LED light sources provide thelarge-field illumination to control cell speed, while the green LED is the light source fordark field microscopy; these light beams pass through a custom-built phase ring above the condenser (see panel B). Excitation light that transmits through the excitation filter below the stage from a Hg lamp is used for wide-field epifluorescence microscopy. Also see Methods. (B) Schematic diagram of the custom-built phase ring. As a key component of the imaging system, it integrates a 3D-printed holder, a flat glass lens, and a shortpass filter (500 nm) that only allows light below a certain wavelength (including blue and violet light) to pass through.

**Fig. S6.**
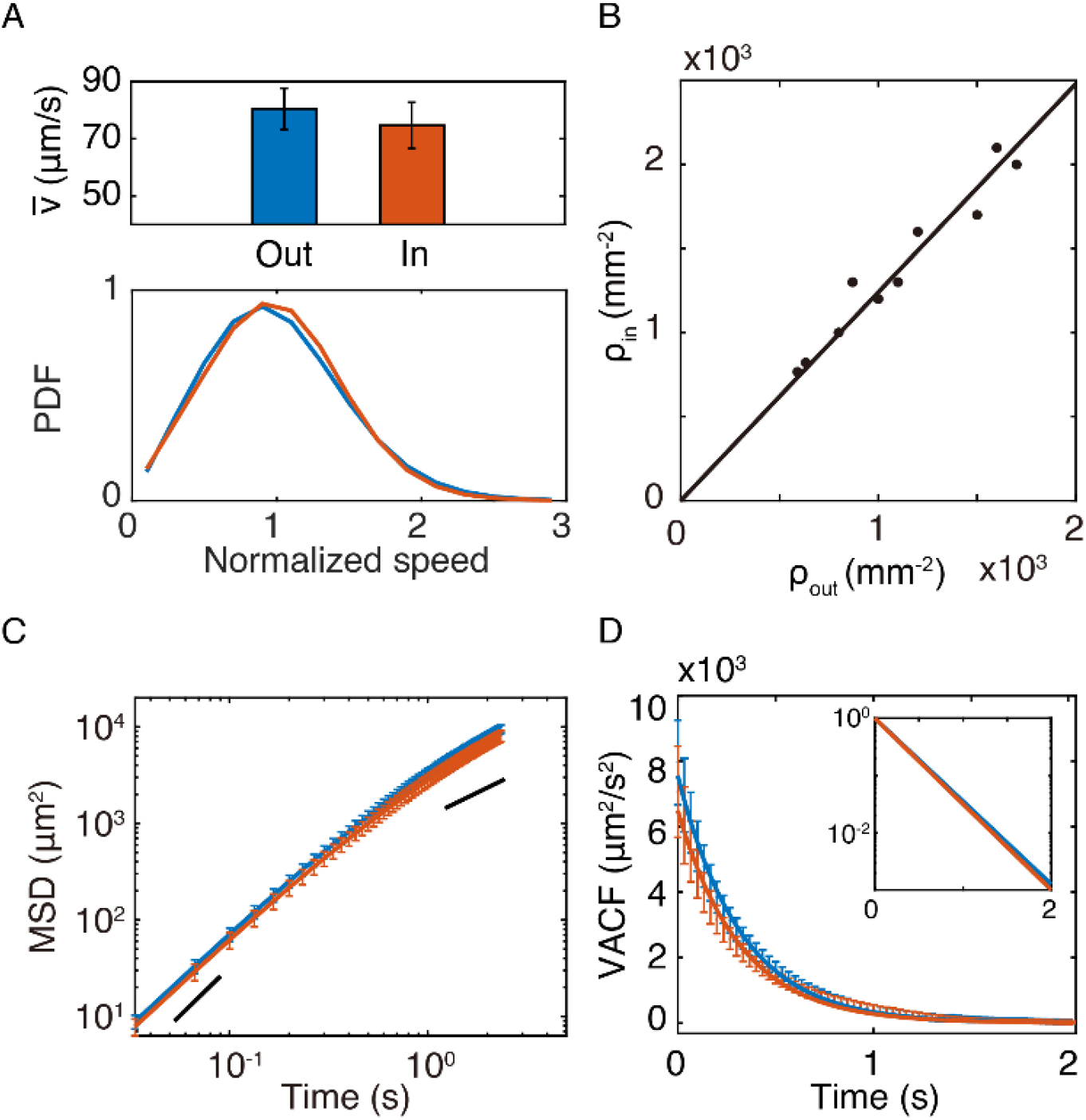
Motion pattern of cells in light-induced liquid bulges. (A) Speed of non-chaining *B. subtilis* Δ*sinI* cells inside and outside the liquid bulges induced by violet light (Methods). Upper panel: Average speed 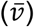 of cells that remained outside and inside the liquid bulges during the tracking period are shown as blue (80.4 ± 7.2 μm/s; mean ± SE, N = 7) and red (74.7 ± 8.0 μm/s; mean ± SE, N = 7) columns, respectively. Lower panel: Average probability density functions (PDFs) of the normalized speed of cells corresponding to the two scenarios in the upper panel, following the same color coding as that of the columns in upper panel. The speed was normalized by the mean speed of all cells in the corresponding scenario. (B) Relation between surface-packing density of fluorescent cells seeded in the swarm inside (*ρ*_*in*_) and outside (*ρ*_*out*_) bulges. Solid line is least-squares linear fit with *ρ*_*in*_=1.*2*4 (R^*2*^= 0.94). Data presented here was from 10 biological replicate experiments. (C) Mean squared displacement (MSD) of cells inside (red) and outside (blue) liquid bulges in logarithmic scale. The lines with slope 1 (diffusive motion) and 2 (ballistic motion) are guides to the eyes. (D) Velocity autocorrelation functions (VACFs) of cells that remained inside and outside the liquid bulges computed based on single cell trajectories. Solid lines are exponential fits to the VACFs [blue: outside liquid bulges, VACF_out_ ∼ exp(-t/0.31), persistence time *τ*_out_ = 0.31 ± 1318 0.01 s; red: inside liquid bulges, VACF_in_ ∼ exp(-t/0.30), persistence time *τ*_in_ = 0.30 ± 0.01 1319 s). The diffusivity computed from VACF is ∼15% lower inside bulges than outside (∼1030 μm_2_/s vs. ∼1210 μm_2_/s) (Methods). Plotted in the inset are normalized fitted VACFs outside (blue) and inside (red) liquid bulges in a semi-logarithmic scale. Error bars represent the standard deviation of 7 biological replicates. The results presented in this figure show that the light-induced liquid bulges are of the same nature as those naturally developed in the swarm.

**Fig. S7.**
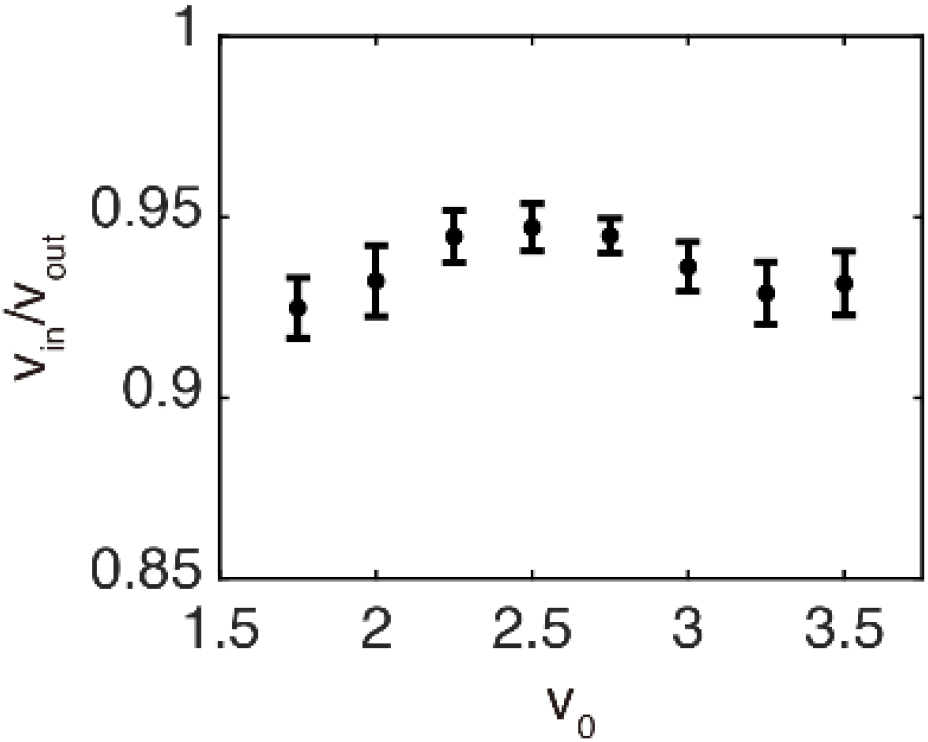
The ratio of apparent particle speed inside and outside bulges for 1329 different particle activity in the active Brownian dynamics simulation. The apparent speeds (*ν*_*in*_ and *ν*_*out*_) were computed for particles during the simulation; the particle activity is denoted by active speed *ν*_0_ set as a constant for all particles outside bulges in a simulation run. Also see Methods. Error bars represent standard deviation (N=10 simulation runs). The ∼5-10% difference in the apparent particle speeds inside and outside bulges is consistent with experimental measurements.

